# Control of atypical PKCι membrane dissociation by tyrosine phosphorylation within a PB1-C1 interdomain interface

**DOI:** 10.1101/2023.01.03.522491

**Authors:** Mathias Cobbaut, Neil Q. McDonald, Peter J. Parker

## Abstract

Atypical PKCs are cell polarity kinases that operate at the plasma membrane where they function within multiple molecular complexes to contribute to the establishment and maintenance of polarity. In contrast to the classical and novel PKCs, atypical PKCs do not respond to diacylglycerol cues to bind the membrane compartment. Until recently it was not clear how aPKCs are recruited; whether aPKCs can directly interact with membranes or whether they are dependent on other protein interactors to do so. Two recent studies identified the pseudo-substrate region and the C1 domain as direct membrane interaction modules, however their relative importance and coupling are unknown. We combined molecular modelling and functional assays to show that the regulatory module of aPKCι, comprising the PB1 pseudo-substrate and C1 domains, forms a cooperative and spatially continuous invariant membrane interaction platform. Furthermore, we show the coordinated orientation of membrane-binding elements within the regulatory module requires a key PB1-C1 interfacial β-strand (BSL). We show this element contains a highly conserved Tyr residue that can be phosphorylated and that negatively regulates the integrity of the regulatory module, leading to membrane release. We thus expose a novel regulatory mechanism of aPKCι membrane binding and release during cell polarization.

## Introduction

Protein kinase C (PKC) isozymes are subject to tight regulation of their catalytic activities rendering them responsive to an array of cellular signals in specific compartments consistent with other AGC kinase superfamily members (1, 2). For classical and novel PKC isoforms, changes in lipid signalling that induce formation of diacylglycerol (DAG) drive activation and therefore PKC responses to various G-protein coupled receptors and receptor tyrosine kinases linked to PLC activity (1). The atypical PKCs (iota and zeta) on the other hand do not contain DAG-responsive elements in their regulatory amino-termini, but instead contain a PB1 domain tethered via a pseudo-substrate linker region to a DAG-unresponsive C1 domain (3). The pseudo-substrate (PS) motif is thought to repress catalytic activity by competitively blocking access to substrates (4), while the PB1 domain mediates protein-protein interactions with activating/regulatory proteins such as p62 and Par6, which in turn are also modular proteins allowing for multivalent protein interactions (5, 6).

Many aPKC functions are intricately linked with its recruitment to the plasma membrane. Indeed, the polarity complexes that define apical and basolateral membrane identity bind membranes specific to these compartments (7). The manner in which aPKC is recruited into and excluded from these membrane-associated polarity complexes is still not entirely clear. Since there is an interdependency with other proteins such as Par3 and Cdc42, membrane association was long considered to be indirect occurring through association with these and other partner proteins (8–11). However two recent studies have shown at that aPKCι can associate directly with cell membranes through its regulatory module, defined hereafter as aPKCι^RM^ (12, 13). The membrane-binding determinants within the aPKCι^RM^ have been identified as the PS region, which contains a stretch of positively charged Arg residues, and the C1 domain. It is unclear hitherto what the relative contribution of each region is. For example, in HEK293 cells, substitutions in the PS region render the protein unable to interact with the plasma membrane, whereas in *Drosophila* neuroblasts the C1 domain has been identified as the dominant membrane interaction module (12, 13).

Here, we approach the question of how multiple aPKC membrane-binding determinants coordinate and how they are coupled. We use Alphafold *in silico* predicted models for the aPKC regulatory module combined with biochemical and cellular assays to validate a continuous surface harbouring known membrane-interacting determinants. We show that the PB1 and C1 domains are coupled and oriented though interfacial contacts also aligning the PS motif and C1 domain for membrane interaction. We identify a highly conserved phospho-Tyr residue within the PB1-C1 interface and show that phosphorylation of this residue modulates membrane-binding capabilities of the regulatory module. We propose that this phosphorylation influences the ability of aPKC to associate with and disassociate from membranes in a regulated manner.

## Experimental procedures

### In silico/molecular modelling

To predict the tertiary structure of the regulatory domain we used Alphafold Colab (https://colab.research.google.com/github/sokrypton/ColabFold/blob/main/AlphaFold2.ipynb) (14) with residues 22-194 of human aPKCι as input. The top scoring model coordinates were used for subsequent prediction of membrane binding motifs using the Membrane Optimal Docking Algorithm (MODA) (15) and ConSurf (16).

### Cell lines and reagents

HEK293T cells were grown in Dulbecco’s modified eagle medium (DMEM) supplemented with 10% (v/v) Fetal Bovine serum (ThermoFisher scientific), 100U/ml Penicillin and 100μg/ml Streptomycin (ThermoFisher scientific). HEK293FS cells were grown in Freestyle 293 Expression Medium (ThermoFisher Scientific). Cells were obtained from the institute’s central catalogue, where they are routinely checked for mycoplasma. The phospho-Tyr136 antibody was made in-house using an immunogenic peptide spanning residues 125-141 of haPKCι with modifications at Tyr-136 (phosphate) and Cys-137 (S-methylcarboxyamide). Anti-thiophosphate ester antibody was from Abcam (Cambridge, UK). Anti-FLAG M2 antibody, and FLAG-agarose resins were purchased from Sigma (St. Louis, MO, USA). Streptactin II agarose was from GE healthcare (Little Chalfont, UK). Myc-trap agarose was from Chromotek (Planegg, Germany) Anti-Myc, secondary HRP-linked goat anti-Rabbit and Horse anti-Mouse antibodies were from Cell Signaling Technologies (Beverly, MA, USA). Polyethyleneimine (PEI) was from Polysciences Inc. (Warrington, PA, USA). Mutagenesis and cloning were done using In-Fusion (Takara, Shiga, JP). Primers and plasmids used are listed in Table S4. Peptides used were made in-house.

### Protein expression and purification

For expression of Strep-tagged Par6 co-expressed with Myc-tagged fl-aPKCι adherent HEK293T cells were transiently transfected using polyethylene-imine (PEI) at a 1:3 (m/m) plasmid/PEI ratio. Forty-eight hours post-transfection, cells were lysed in 50 mM Tris, pH 7.4, 150 mM NaCl, 1% Triton, 0.5mM TCEP supplemented with phosphatase inhibitors (Phosphostop, Roche, Germany), and protease inhibitors (cOmplete, Roche, Germany). Cell lysates were incubated with Streptactin II (GE healthcare) affinity beads for 2 hours at 4°C while rotating. Next, the beads were washed twice with Lysis buffer supplemented with NaCl to 500mM, once with Lysis buffer and once with elution buffer (50mM Tris pH7.4, 150mM NaCl, 0.5mM TCEP). The protein was eluted in elution buffer supplemented with 2.5mM D-desthiobiotin. For the purification of 2xStrep-tagged fl-aPKCι the same protocol was followed but HEK293FS cells were used and transfected for 96h instead of 48h. In addition, a size exclusion chromatography step was performed on a Superdex-200 Increase column and peak fractions were pooled, dialyzed against elution buffer and concentrated. Protein quantity and purity was assessed using SDS-PAGE and A280.

### *In vitro* kinase assay

HEK293T cells expressing myc-tagged fl-aPKCι WT and mutants were lysed in lysis buffer (50 mM Tris, pH 7.4, 150 mM NaCl, 1% Triton, 0.5mM TCEP supplemented with phosphatase inhibitors (Phosphostop, Roche, Germany), and protease inhibitors (cOmplete, Roche, Germany)) and immunoprecipitated using Myc-Trap agarose (Chromotek). Beads were washed once in lysis buffer containing 500mM NaCl and twice in TBS. Equal amounts of the bead suspension were then added to 1.5μg myelin basic protein (MBP) and 250μM ATP-γS in assay buffer (1x final; 50mM Tris, 10mM MgCl_2_). Reactions were incubated at 30°C for 30 min and subsequently added with 2.5 mM p-Nitrobenzyl mesylate (PNBM) for 1.5h at room temperature. Incorporation of S-labeled phosphates was followed via western blotting with an anti-thiophosphate ester antibody.

### Differential scanning fluorometry

Protein at 250μg/ml was incubated with SPYRO-orange protein dye (1:400) in 50mM Tris, 150mM NaCl, 0.5mM TCEP with or without 1mM ATP and 10mM MgCl_2_. The reactions were preincubated at room temperature for 5 min before being distributed into a MicroAmp 96-well qPCR plate. Samples were heated from 25-98°C in a QuantStudio Flex 12 RT-PCR machine (ThermoFisher Scientific). Measurements of fluorescence emission at 570nm were followed for each 0.3°C increment. Raw curve data was exported and derivative curves were plotted. The T_m_ of protein in each well was determined as the temperature at the inflection point of the melting curve using the maximal point of the derivative.

### Immunofluorescence

Cells were grown on 13□mm glass coverslips, transfected with constructs as indicated and 24h post transfection cultures were treated with 500nM nocodazole for 16h before fixation or left untreated. Cells were fixed with 4% PFA and permeabilized with PBS+0.1% Triton X-100. Coverslips were then blocked in 3% BSA in PBS and incubated with FLAG M2 antibody (1:500). Secondary antibody was 1:1500 goat anti-mouse 488 (Thermo Fisher, A11001). Coverslips were mounted using Prolong gold with DAPI (Thermo Fisher Scientific) and imaged. All the images were acquired using an inverted laser scanning confocal microscope (Carl Zeiss LSM 880) using a 63x Plan-APOCHROMAT DIC oil-immersion objective. The same exposure settings were used to acquire all images (excluding DAPI channel). Images shown in figures were processed in ZEN Blue edition (Zeiss) where a gold LUT pseudo-colour transformation was applied to the 488-emission channel. All images were batch-processed to adjust brightness/contrast. Quantification of membrane and cytoplasmic intensities was done on the raw images in ImageJ. The whole membrane region was traced with the freehand tool and the average pixel intensity was recorded. In the same way the average intensity for a trace inside in a sub-membrane region of the cytoplasm was recorded and the fraction of recorded intensities was plotted.

### Cell fractionation

To assess phosphorylation of Tyr-136 in the membrane and cytosolic fractions, HEK293T cells expressing FLAG-tagged regulatory domain were fractionated as in (17). Briefly, cells were lysed in Homogenization buffer (20 mM Tris, pH 7.4, 10mM EDTA, supplemented with phosphatase inhibitors (Phosphostop, Roche, Germany), and protease inhibitors (cOmplete, Roche, Germany)) and subjected to sonication. The suspension was then centrifuged at 100.000xg for 20’ at 4°C and the supernatant (Soluble fraction) was collected. The pellet (Particulate fraction) was resuspended in homogenization buffer containing 1% Triton X-100, sonicated and again centrifuged at 100.000g for 20’ at 4°C. The supernatant was collected and FLAG-IP was performed on both supernatants to isolate the regulatory domain for Western analysis with the pY136 antibody.

### Reduction-alkylation for immunodetection with pTyr-136 antibody

Western blots analysing levels of Tyr-136 phosphorylation were performed according to standard procedures but with the subsequent additional steps. After transfer the nitrocellulose membrane was incubated overnight in PBS +10mM DDT. It then was washed twice in PBS and incubated with PBS +10mM iodoacetamide (Sigma) for 1h at room temperature. The membrane was washed twice with PBS and immunoblotting was continued according to standard procedures.

### Lipid overlay assay

aPKCι-Par6 WT and mutant complexes were purified as described above using 2xStrep-Par6 as bait. The protein complexes were analysed via silver staining and incubated with lipid coated membranes (PIP-Strips cat. P-6001) (Echelon Bioscience). The PIP strips were subsequently probed with Myc-antibody (1:1000) and HRP-tagged secondary antibody (1:3000). Membranes were processed and exposed in parallel in an Imagequant LAS 4000 (GE Healthcare).

## Results

### A continuous membrane-binding surface is oriented within a compact PB1-PS-C1 aPKCι regulatory module

In light of the recent discrepancies in the relative contributions of PS and C1 domain to aPKC membrane interaction, we took a holistic approach to understand whether the regulatory module acts as a functional unit contributing two coupled membrane-binding determinants. We used AlphaFold Colab to predict the aPKCι regulatory module (aPKCι^RM^) spanning the PB1 domain, the PS motif and the C1 domain (14). The top five structural models predict a compact, globular shape with buried inter-domain contacts suggesting a coupled functional unit (Fig S1A). The models predict that the PB1 and C1 domains are crucially bridged together via an interdomain β-strand which we term beta-strand linker (BSL) (Fig 1A), that is part of a continuous sheet with the β-strands in the PB1 and C1 domains with a parallel orientation to the PB1 and antiparallel to the C1 domain. All of the models show very low predicted alignment errors (PAE), indicative of high confidence in the continuous fold (Fig S1B). Using the highest scoring AF Colab model as input, we set out to predict membrane binding residues via the Membrane Optimal Docking Algorithm (MODA) package (15). Both the PS and two regions in the C1 domain (termed loop 1 and loop 2) have high membrane-association prediction scores (Fig 1B,C, Table S1). Interestingly, both regions form a continuous surface containing both positively charged and hydrophobic residues (Fig 1C,D), potentially contributing to a single composite membrane interaction site. The importance of these residues is further underscored by the fact that most residues constituting the platform are highly conserved as shown via ConSurf analysis (Fig S1E, Table S2), while there is less stringency for amino acid composition on the opposite side of the domain (Fig S1F).

**Figure 1.**
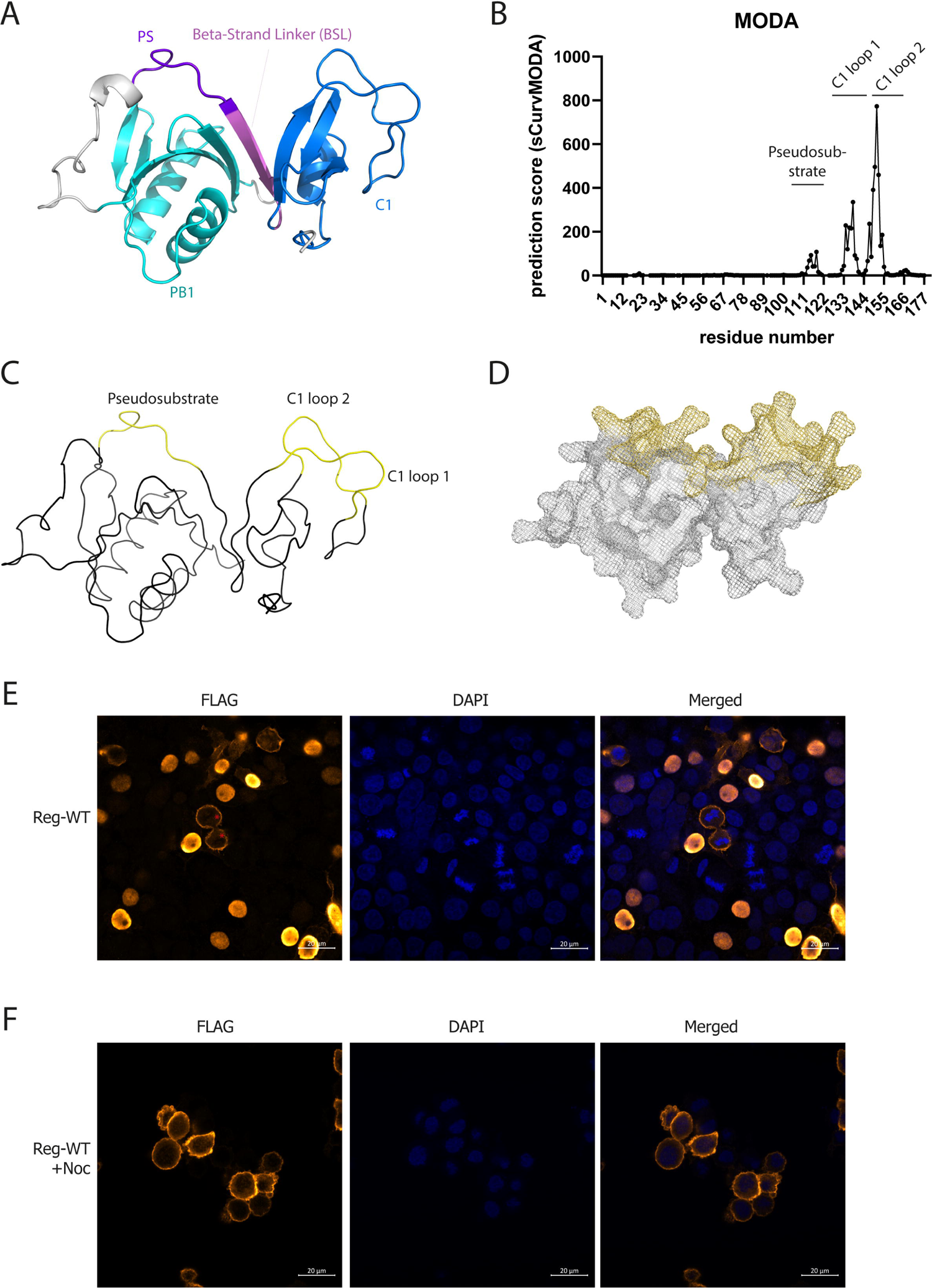
A. Representation of the aPKCι regulatory module as predicted by Alphafold Colab (best scoring model). The individual domains and regions are indicated and coloured; from N-C PB1 domain (cyan), Pseudo-substrate (PS) region (dark purple), Beta-Strand Linker (BSL) (light purple), C1 domain (blue). B. Per residue prediction score of the MODA prediction with above-threshold scoring regions indicated. C. Cartoon diagram (omitting secondary structure features) of predicted membrane interacting regions from the MODA prediction displayed in the AF Colab model. D. Mesh representation of panel C showing a continuous sidechain surface E. Localization of FLAG-tagged aPKCι regulatory module in HEK293T cells; red asterisks indicate mitotic cells. F. Localization of FLAG-tagged aPKCι regulatory module in nocodazole-treated (500nM, 16h) HEK293T cells.

To probe the intrinsic capabilities of aPKCι^RM^ in membrane binding, we expressed FLAG-tagged aPKCι^RM^ in HEK293T cells. We observed a striking localization pattern that was cell cycle-dependent. In interphase cells the regulatory module is predominantly localized in the nucleus (Fig 1E), whereas in mitotic cells, aPKCι^RM^ is uniformly associated with the plasma membrane (Fig 1E, asterisks). This observation is in line with those from *Drosophila* neuroblasts where over-expressed aPKCι^RM^ constructs display similar behaviour in interphase and at mitosis (13). This property allowed us to assess the membrane binding requirements of aPKCι^RM^ in a physiological context by arresting cells in mitosis with nocodazole (Fig 1F). To exploit this “mitotic trap” assay, we introduced site-specific mutations targeting residues with the highest relative prediction scores for membrane interaction (Table S1). While the WT regulatory module has a strong (∼4-fold) enrichment at the plasma membrane in nocodazole treated cells (Fig 2A), we observed a complete loss of membrane binding when substituting the highly predictive ^160^RIWGL^164^ sequence in the C1 domain (loop 2) with ^160^AIAGD^164^ (Fig 2B,F). Similarly, a complete loss of membrane binding was observed with ^126^RRGARR^131^ to ^126^AAGAAA^131^ substitutions in the PS region (termed PS 4R.A, Fig 2C,F). Substitution of Arg-147 or Arg-150/151 to Ala in the C1 domain (loop 1) had a smaller impact reducing membrane binding by about 1.5-fold (Fig 2D,E,F). This indicates that both regions in the C1 domain and the PS motif contribute to allow optimal membrane binding in these mitotic cells. Based upon the model, it is likely these motifs are organized in a continuous surface, the integrity of which likely dictates the capacity to bind membranes.

**Figure 2.**
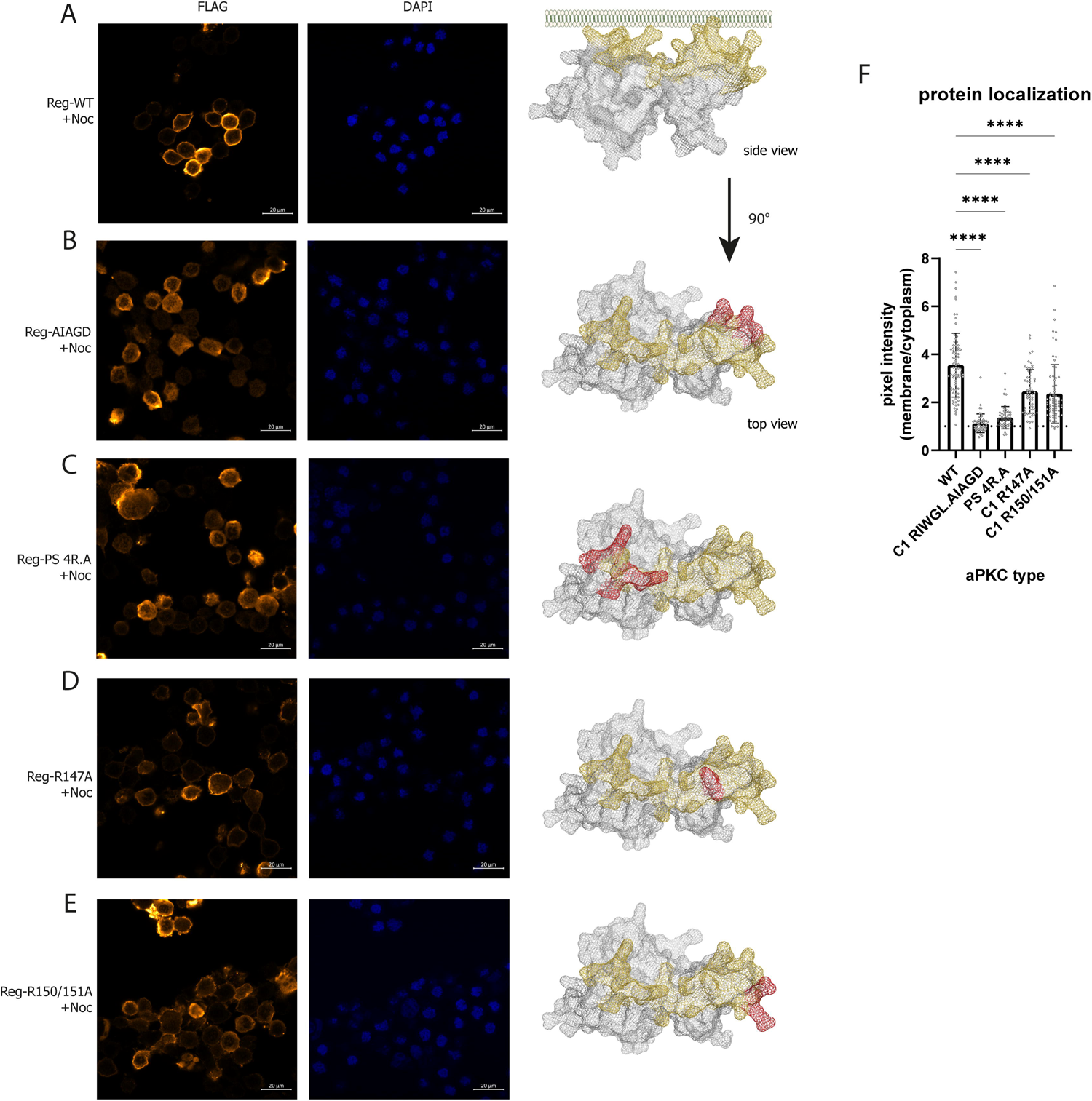
A. Localization of WT FLAG-tagged regulatory module in nocodazole-treated (500nM, 16h) HEK293T cells. B-D. Expression of the indicated mutant FLAG-tagged regulatory modules in nocodazole-treated (500nM, 16h) HEK293T cells. Mutated residues are depicted on the protein model in red. E. Quantification of membrane/cytoplasmic pixel intensities measured in >45 cells imaged via confocal microscopy. Representative experiment of 3 biological replicates. Differences analysed via one-way ANOVA (****P ≤ 0.0001)

### Tyr-136 phosphorylation within an interfacial motif uncouples membrane-binding determinants but does not impact catalytic activity

One of the more salient features of the regulatory module is the interfacial BSL motif positioning the PB1 and C1 domain, orienting the membrane-binding determinants within the aPKCι^RM^. The BSL shows overall high conservation centred on a centrally positioned, highly conserved Tyr residue 136 (Fig S1G, Table S2). Inspection of Phosphosite indicated that this residue is frequently found to be phosphorylated in more than 60 high-throughput studies in a variety of conditions and cell types, alongside the familiar activation loop and turn motif priming sites (Fig 3A) (18). Consistent with this, it is also one of few aPKCι phospho-residues identified in the MaxQuant Database (MaxQB) (Table S3) (19).

**Figure 3.**
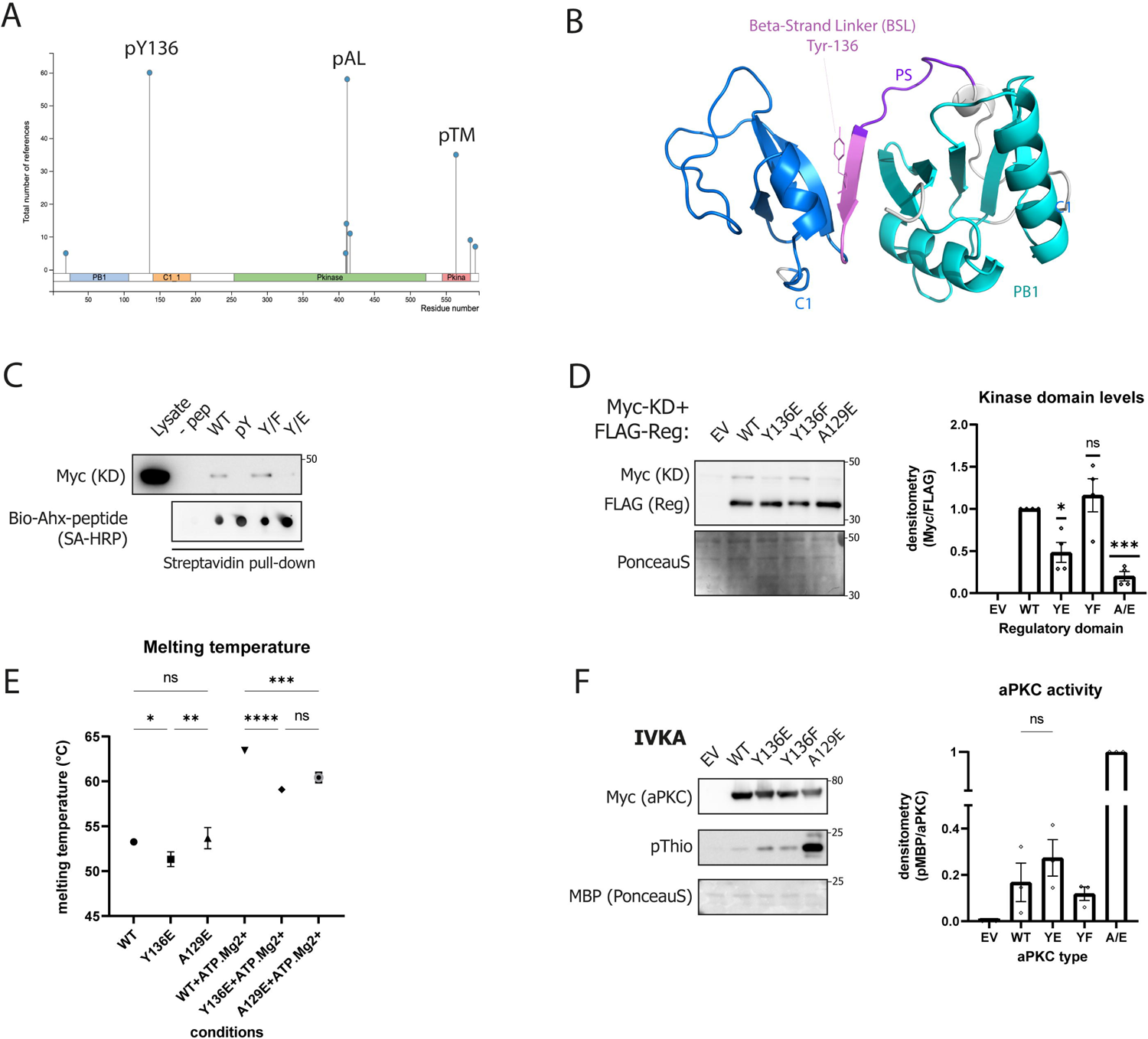
A. Identification of main phosphorylation sites on aPKCι identified in the Phosphosite database B. visualization of Tyr-136 central to the BSL region of the regulatory module, colour coded as in Fig. 1A C. Peptide-pulldown of Myc-tagged aPKCι kinase domain with extended pseudosubstrate peptides (Biotin-Ahx-SIYRRGARRWRKLYCANGHT-CONH2) containing the indicated modifications at Y^14^, i.e. pY, Y/E and Y/F. D. Co-expression of FLAG-tagged regulatory module and Myc-tagged kinase domain of aPKC in HEK293T cells. Equal amounts of DNA were transfected and stabilization of Myc-tagged kinase domain by different mutants of the regulatory module was followed. Quantification of Myc-kinase domain levels relative to WT aPKCι^RM^ co-expression shown in the right panel (n=3 biological replicates, average +/-SEM, analysed via one-sample t-test (v.a.v. a value of 1): ns P > 0.05, * P ≤ 0.05, *** P ≤ 0.001). E. DSF assay measuring thermal stability of WT and mutant fl-aPKCι in absence or presence of ATP.Mg^2+^. Plotted is the T_m_ derived from the inflection point of the melting curve (using the derivative maximum). F. *In vitro* kinase assay (IVKA) with immunoprecipitated WT or mutant kinases on MBP. Incorporation of new phosphates into MBP was followed with ATP-γS. Quantification of MBP thiophosphate levels shown on the right (n=3 biological replicates, average +/-SEM, analysed via unpaired t-test: ns P > 0.05)

As Tyr-136 lies at the core of the regulatory module within the interfacial BSL motif and proximal to the PS (Fig 3B), we considered whether tyrosine phosphorylation could influence the full-length kinase conformation, stability and activation state. First, we probed whether phosphorylation affects the interaction between the PS region and the kinase domain. By definition, the PS is proposed to bind the catalytic domain at the substrate-binding pocket to be released upon aPKCι activation. We wondered whether in a linearized context, Tyr phosphorylation proximal to the PS could influence this behaviour. To test this, we immunoprecipitated a Myc-tagged kinase domain (aPKCι^KD^) from cell lysates using a biotin-labeled extended pseudo-substrate peptide (ePSP) containing either a non-phosphorylated Tyr or pTyr residue or Glu or Phe substitutions to mimic either phosphorylated or non-phosphorylated states respectively (Fig 3C). We saw specific binding of the peptide to the core kinase domain and observed loss of kinase domain interaction with the peptide when the Tyr residue is either phosphorylated or substituted with Glu. Conversely a non-phosphorylated or Phe substituted peptide effectively captured the aPKCι^KD^. To confirm whether the phosphorylation of this residue reduces the interaction between aPKCι kinase domain and its regulatory module, we assessed the protection of the kinase core by co-expressing the isolated kinase domain with either wild type (WT) or mutant forms of aPKCι^RM^ (Fig 3D). While the WT aPKCι^RM^ protein product protects the aPKCι^KD^, a Y136E substitution causes a reduction in the levels of aPKCι^KD^ protein, as does a previously described pseudo-substrate disengaged A129E mutant (4). These results indicate that the kinase domain-regulatory module interaction is weakened by phosphorylation of Tyr-136.

To probe whether this effect is due to loss of direct binding of the PS region to the kinase domain or the aPKCι^RM^ conformation, we purified WT, Y136E and A129E mutant full-length aPKCι (aPKCι^FL^) from HEK293FS cells and subjected them to differential scanning fluorometry (DSF) (Fig 3E). In the apo-form, the melting temperature (T_m_) for the WT protein is 53.3°C, similar to the A129E mutant (T_m_ = 53.7°C). The Y136E mutant is less thermostable with a T_m_ of 51.3°C in the apo-form. Addition of ATP.Mg^2+^ stabilizes both WT and mutant kinases, however comparatively the A129E mutant is stabilized to a lesser degree than WT (ΔT_m_= 6.7°C vs WT ΔT_m_= 10.2°C). This suggests that occupation of the active site with ATP, while stabilizing the kinase core, may result in a less thermostable open conformation, possibly because of phosphate-driven charge repulsion with the Glu substitution. The Y136E mutant is less thermostable than WT in both the presence and absence of ATP, indicating that decreased thermostability is caused by mutation-driven alterations in the regulatory module.

To test the impact of Tyr-136 phosphorylation on aPKCι catalytic activity, we followed the phosphorylation of MBP *in vitro* after immunoprecipitation of WT and mutant forms of aPKCι^FL^. To monitor the incorporation of new phosphate groups on MBP we made use of γ-thiophosphate ATP (Fig 3F). MBP was readily phosphorylated by the conformationally “open” A126E mutant but showed only a marginal nonsignificant increase in phosphorylation upon incubation with a Y136E mutant compared to WT. This indicates that in the ‘tethered’ state of the full-length protein, the impact of the weakened regulatory module/catalytic domain interaction consequent to Tyr-136 phosphorylation is modest compared to the pseudo-substrate A129E mutation, while there is a strong effect on the overall aPKCι^RM^ conformation.

### Tyr-136 phosphorylation disrupts aPKCi^RM^ membrane binding in cells

Since the above results indicate a change in regulatory module conformation with little impact on activity, we tested whether Tyr-136 phosphorylation had a direct influence on the aPKCι^RM^ membrane binding properties. We assessed membrane occupation in mitotic cells as above, expressing the WT regulatory module (Fig 4A) or mutants in nocodazole treated cells; the PS 4R.A mutant was used as a control for loss of membrane binding. Substitution of Tyr-136 with Glu resulted in a significant 1.5-fold reduction of membrane intensity (Fig 4B,F), whereas a Tyr-Phe substitution had a negligible impact on membrane binding (Fig 4C,F), consistent with Tyr phosphorylation being disruptive for membrane interaction. Interestingly a substitution of A129 to Glu had only a small but significant impact on membrane binding capabilities, despite its negative charge central to the PS region (Fig 4D,F), whereas substitutions of the PS Arg residues resulted in a complete loss of membrane binding (Fig 4E,F).

**Figure 4.**
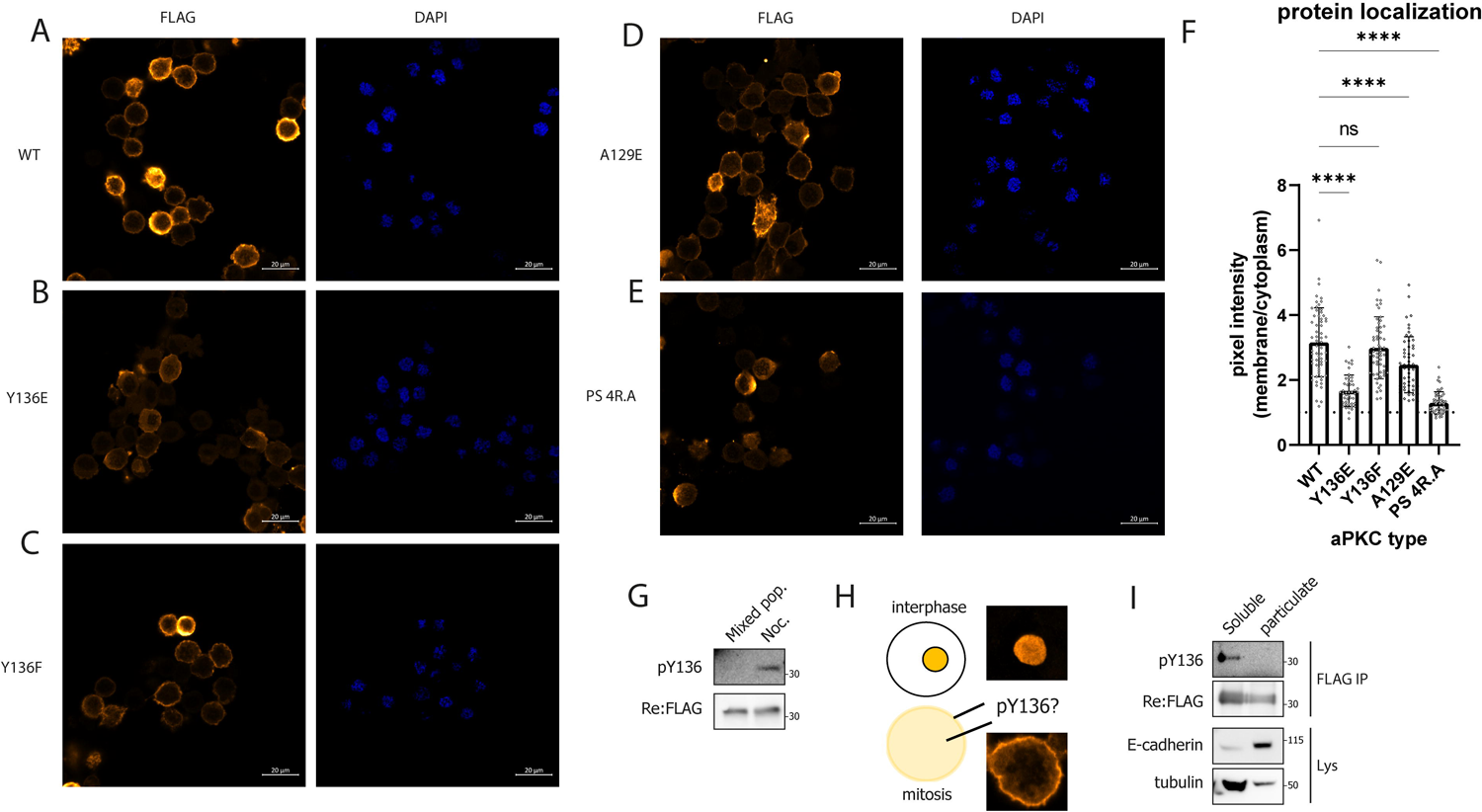
A-E. Expression of the indicated mutant FLAG-tagged regulatory modules in nocodazole-treated (500nM, 16h) HEK293T cells. F. Quantification of membrane/cytoplasmic pixel intensities measured in >45 cells imaged via confocal microscopy. Representative experiment of 3 biological replicates. Differences analysed via one-way ANOVA (ns P > 0.05, ****P ≤ 0.0001) G. Western blot analysis of phospho-Tyr-136 levels in the in untreated (mixed pop.) cells or cells treated with 500nM Nocodazole for 16h (Noc.). H. Representation of the observed localization of regulatory module species and possible mapping of its phospho-form. I. Fractionation of cells by ultracentrifugation and probing for Tyr-136 phosphorylation of the regulatory module (representative experiment of n=2 biological replicates).

To probe Tyr-136 phosphorylation in the regulatory module of aPKC we developed a site-specific and phospho-specific rabbit antibody. Initial attempts resulted in poor and nonspecific immunoreactivity, but using a derivatized phosphopeptide with S-methylcarboxyamide coupled to the adjacent Cys residue C-terminal to the phosphorylated Tyr residue, we were able to detect Tyr phosphorylated Myc-aPKC in pervanadate treated cells after immunoprecipitation (Fig S2A). The alkylation greatly improved epitope recognition, specific to nitrocellulose membranes following alkylation of proteins with iodoacetamide, confirming the double-modified epitope recognition (Fig S2A,B). Interestingly, using this antibody, we could show significant levels of Tyr-136 phosphorylation in regulatory module constructs in nocodazole treated cells (Fig 4G). The Y136 phosphorylation of the ‘free’ aPKCι^RM^ in mitotic cells, suggested that this was not due to the aPKCι kinase domain exhibiting Tyr specificity due to proximity (*in casu* the PS motif), as seen in other Ser/Thr kinases as recently observed in the PKDs (21). Indeed, we could detect no phosphorylation of Tyr-136 upon incubation of aPKCι^FL^ with ATP.Mg^2+^ (data not shown). Exploring candidate upstream kinases, we pulled down proteins from a cell extract using a biotinylated ePSP peptide containing either a Y, F or pY residue corresponding to Tyr-136 and then spiked the precipitated protein mixture with ATP.Mg^2+^, allowing any associated kinase to (auto)phosphorylate. Western blot analysis with a phospho-Tyr antibody showed tyrosine kinase (autophosphorylation) activity around ∼60kDa and ∼170kDa in protein mixtures associating with the Y and F but not the product pY peptide (Fig S3A). In a subsequent screen with a panel of Tyr kinase inhibitors we found responsiveness of aPKC ^RM^ Tyr-136 phosphorylation in nocodazole treated conditions to the Src family kinase (SFK) inhibitors PP2 and dasatinib (Fig S3B). This implicates SFKs in targeting this site and is consistent with the finding of a ∼60KDa tyrosine phosphorylated protein(s) captured on the site-affinity matrix.

The fact that Tyr-136 is phosphorylated in nocodazole treated cells led us to ask the question whether the phosphorylated species segregates to the membrane or cytosolic compartment in mitotic cells (Fig 4H). To assess any preferential localization of the phosphorylated species we fractionated nocodazole-treated cells by ultracentrifugation (100,000xg) to separate membrane (particulate) and cytosolic (soluble) components. We could detect no phosphorylation of the regulatory module in the particulate fraction indicating that this species does not associate with the membrane fraction, while phosphorylated protein was detected in the cytosol (Fig 4I). This further supports our observation with the Glu phospho-mimetic mutant, showing that tyrosine phosphorylation impacts the protein cycling between membrane and cytosolic fractions.

To further assess direct lipid binding in the context of the full-length protein, including the effects of increased dynamics between regulatory and kinase domain in the mutants, we purified the aPKC-Par6 complex from HEK293T cells using Par6 as bait (Fig 5A). The binary complex is thought to be in a more open conformation as Par6 is proposed to displace the PS region, potentially exposing the aPKCι^RM^ for lipid binding (4). These binary protein complexes were then used in a lipid overlay assay with PIP-strips (Fig 5B). The WT complex interacts preferentially with monophosphorylated inositol headgroups, (PI3P, PI4P, PI5P) but also with doubly phosphorylated headgroups (PI(3,4)P_2_, PI(3,5)P_2_,PI(4,5)P_2_) with no strong preference for the position of the phosphate (Fig 5B). The complex also binds phosphatidylserine while we observed no binding to Sphingosine-1-phosphate (S1P) which was previously described as a lipid activator for aPKCs (20). Substituting Tyr-136 with Glu results in reduced binding to all phospholipids, whereas a Phe mutant displays WT characteristics (Fig 5B). Interestingly a PS-disengaged mutant (A129E) results in increased lipid binding despite the glutamic substitution. This is indicative of the fact that this mutation can further drive the Par6-bound open conformation, allowing for enhanced lipid binding. Consistent with the mitotic trap assay the Ala to Glu substitution only seems to have a small effect on membrane binding.

**Figure 5.**
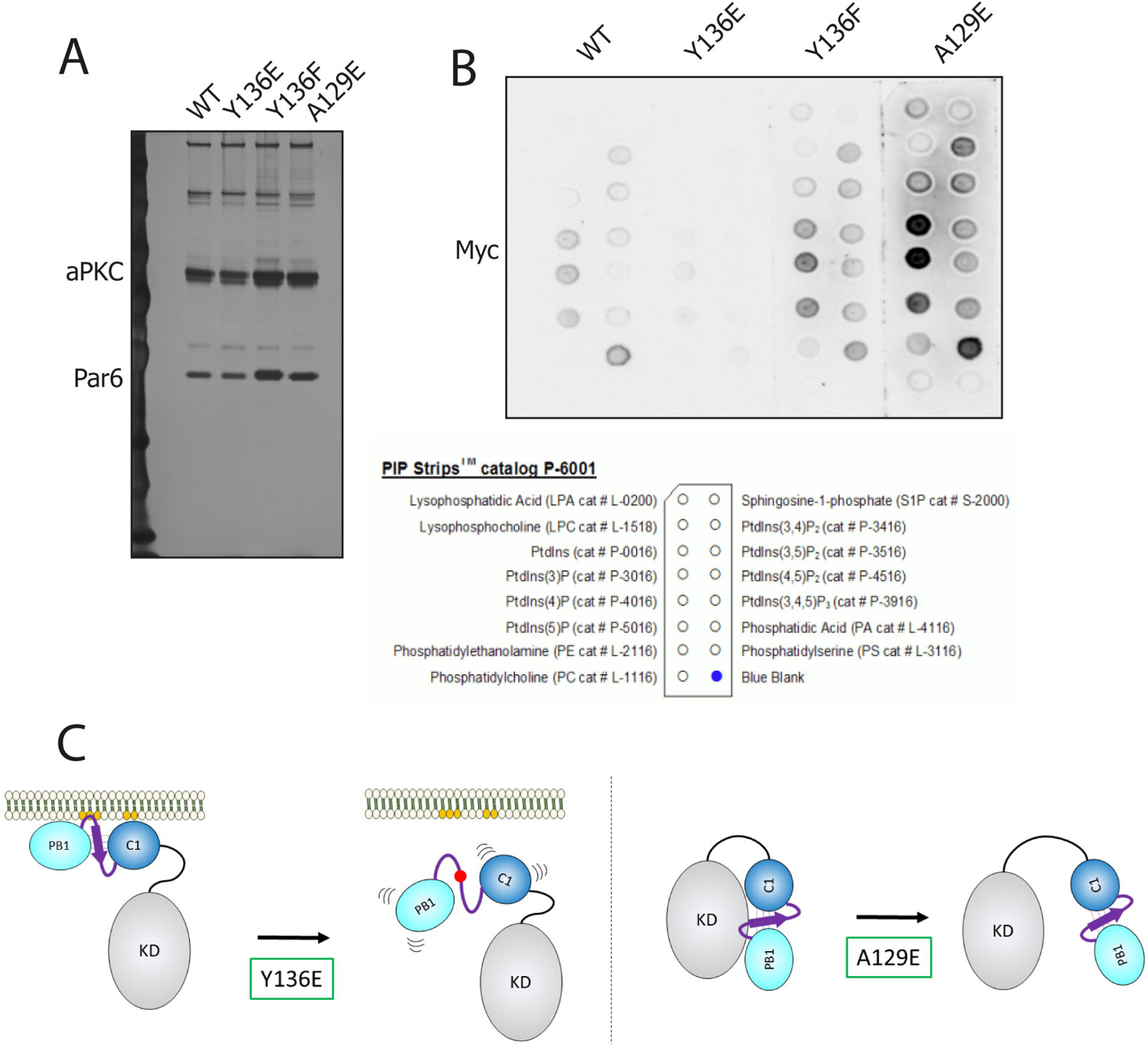
A. Silver-stained SDS-PAGE gel of the Myc-aPKC-2xStrep-Par6 binary complexes purified through 2xStrep-Par6. B. Lipid overlay assay using the protein complexes defined in A. Levels of aPKCι bound to the PIP-strip membrane were quantified using anti-Myc tag antibody. C. schematic representation of the impact of mutants and phospho-states of aPKCι. The left panel displays the anticipated consequence of Tyr-136 phosphorylation in the BSL as mimicked by a Y136E substitution, disrupting the continuous membrane binding platform constituted by the PS and C1 domain. The right panel displays the impact of a pseudo-substrate disengaged conformation, mimicked by an A129E substitution which mainly results in relieved auto-inhibition and increased catalysis.

## Discussion

Cell polarity - the spatially asymmetric organization of a cell and its components - is an essential biological process, orchestrated by key regulatory inputs. One central component of this process conserved in higher eukaryotes is atypical protein kinase Cι. Together with its partner Par6 this kinase is involved in regulating a variety of protein complexes, prominently those including Par3, Lgl1/2, Crb, but also Dlg, Par-1, FARP2, Yurt, and others (22–28). While there are functional data on the interdependency of polarity components, molecular insight into the regulation of these assemblies and their distribution is still lacking. For example, it is not clear how the aPKCs are recruited to membranes and membrane-associated protein assemblies. Several studies have shown a dependency on Par-3 (8–11), however, Par3 independent mechanisms have also been described (29). The correct lipid environment plays a crucial role in recruiting and tethering the right polarity components, with anionic phospholipids and prominently PI(4,5)P_2_ playing a dominant role in the apical domain (30, 30, 31). Several polarity proteins respond to these lipids via basic-hydrophobic (BH) motifs (32, 33). Interestingly, in a recent study the pseudo-substrate of aPKCs was identified as such a lipid-responsive BH domain in cells (12). This aligns with the long-standing notion that the PKC pseudo-substrate region interacts with negatively charged lipids (34). In addition the response to PIP3 downstream of insulin has also been mapped previously to the PS region (35). However a subsequent study by the Prehoda group showed that in dividing drosophila neuroblasts membrane targeting was dependent on the C1 domain, also via an anionic lipid-binding mechanism (13).

Here we have investigated membrane binding by aPKCι^RM^ making use of recent advances in AI by Alphafold to derive a structural model for multi-domain modules of proteins. While attempts at purification of aPKCι^RM^ for structure determination were unsuccessful, AlphaFold Colab predictions indicated it adopts a single compact unit comprising the PB1 and C1 domains, in which they are tethered via an interdomain β-strand proximal to the pseudo-substrate embedded in a continuous sheet spanning across both domains. Based on this predicted model we used the MODA prediction algorithm which identified 3 regions containing putative membrane-binding residues. Interestingly, together these predicted membrane-binding regions form a highly conserved continuous platform in the aPKCι^RM^ model. Aside from the PS sequence ^126^RRGARRWRK^134^ we identify ^160^RIWGLGRQ^167^ in the C1 domain as a second sequence strongly contributing to membrane binding. Mutagenesis in either of these sequences completely disrupts membrane binding in the mitotic cell assay, indicating that neither alone provides sufficient affinity and an intact platform is required to establish the avidity effect for optimal membrane binding. Additionally, Arg-147 and Arg-150/151 in the C1 domain also contribute albeit to a lesser degree.

The organization of the membrane binding platform is crucially driven by the interfacial BSL motif, with at its core an evolutionarily conserved phospho-Tyr residue that allows for regulatory input. Interestingly we observed this dynamic behaviour in mitotic cells where we found that aPKCι Tyr-136 was phosphorylated only in the cytoplasm. Whether the released protein retains an active, open conformation impacting cytosolic targets has yet to be determined, although our evidence indicates that a tyrosine phosphorylation mimetic alone is not substantially activating despite weakening the inhibitory interaction of the regulatory module on the catalytic domain (Table 1). Based on the effects on thermostability, we propose that the PB1 and C1 domains become discontinuous when Tyr-136 in the BSL is phosphorylated, compromising the overall conformation and membrane binding (Table1, Fig 5C, left hand side). This distinguishes the BSL from the pseudo-substrate, which, besides being part of the aPKC membrane interaction module, regulates kinase activity by direct interaction with the kinase domain evidenced by a loss of kinase domain stabilization and increased catalytic activity upon A129E substitution (Table 1). The pseudo-substrate is therefore the main regulatory unit influencing catalysis (Fig 5C, right hand side).

**Table 1.**
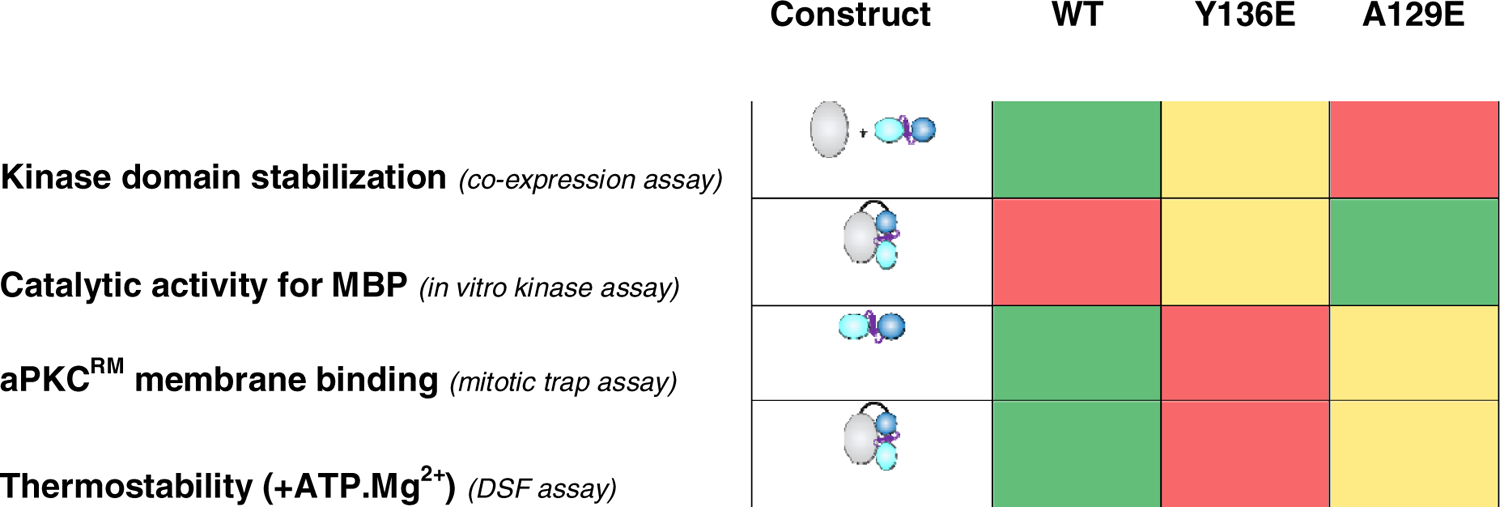
effects measured in the respective assays. Colour-coded based on the average measurements obtained in the assays, cells conditionally formatted from red (low) to green (high).

|  | Construct | WT | Y136E | A129E |
| --- | --- | --- | --- | --- |
| **Kinase domain stabilization** (*co-expression assay*) |  | Green | Yellow | Red |
| **Catalytic activity for MBP** (*in vitro kinase assay*) |  | Red | Yellow | Green |
| **aPKC<sup>RM</sup> membrane binding** (*mitotic trap assay*) |  | Green | Red | Yellow |
| **Thermostability (+ATP.Mg<sup>2+</sup>)** (*DSF assay*) |  | Green | Red | Yellow |

The observation that there is a pTyr residue central to the regulation of membrane binding implicates cross-talk with Tyr kinase activities contributing to a polarized distribution of the aPKCs. The role for Tyr kinase activity in junction formation and cell polarity was clear early on (36–38), however, few molecular links have been established. We show for Tyr-136 that SFKs are potential upstream regulators. Interestingly, Src has long been known to associate with aPKCs, docking to their PXXP rich motifs N-terminal to the pseudo-substrate site; an in vitro phosphorylation reaction with Src using aPKC-derived peptides also included a peptide containing pTyr-136, further supporting the fact that SFKs can act upstream (39, 40). The cell-cycle dependency of Tyr-136 phosphorylation we observed raises intriguing questions for further exploration regarding the Src-aPKC connection in asymmetrical cell division. v-Src transformation of the zebrafish embryo enveloping layer epithelium causes mitotic-dependent extrusion, partially by affecting cell polarity through Cdc42 and aPKC (41 42). Src also promotes epithelial polarity switching in colorectal cancer spheroids, promoting a switch from apical-in (central lumen) to apical-out oriented metastatic cell clusters (43). Other Src family kinases such as Fyn and Lyn have also been involved in the regulation of cell polarity, opening up further possibilities of connecting PTK and polarity pathways (44, 45).

Recruitment and retainment of aPKC at membranes involves a variety of factors that are either PKC intrinsic or extrinsic through membrane-anchored proteins. This complex mesh allows for a variety of inputs and affinity thresholds. We show here the mechanism by which aPKCs can interact with membranes through a continuous interaction platform spanning the whole regulatory region, orchestrated by an interdomain β-strand that can be regulated via tyrosine phosphorylation. The hierarchy and local mechanisms by which aPKCs are recruited and retained at the correct membrane locus, as well as a further understanding of tyrosine kinase crosstalk with cell polarity, are intriguing questions for future exploration.

## Acknowledgements

We thank the Crick Microscopy, Structural Biology and Peptide Synthesis science technology platforms for reagents and expertise. We thank Christopher Earl and David C. Briggs for crystallization efforts and lab members in the Parker and McDonald labs for constructive discussions. The study was funded by the Francis Crick Institute, which receives its core funding [FC001115 and FC001130] from Cancer Research UK, UK Medical Research Council and the Wellcome Trust.

