## Supplemental Data for "Control of atypical PKCι membrane dissociation by tyrosine phosphorylation within a PB1-C1 interdomain interface"

### Supplementary information

#### Figure S1

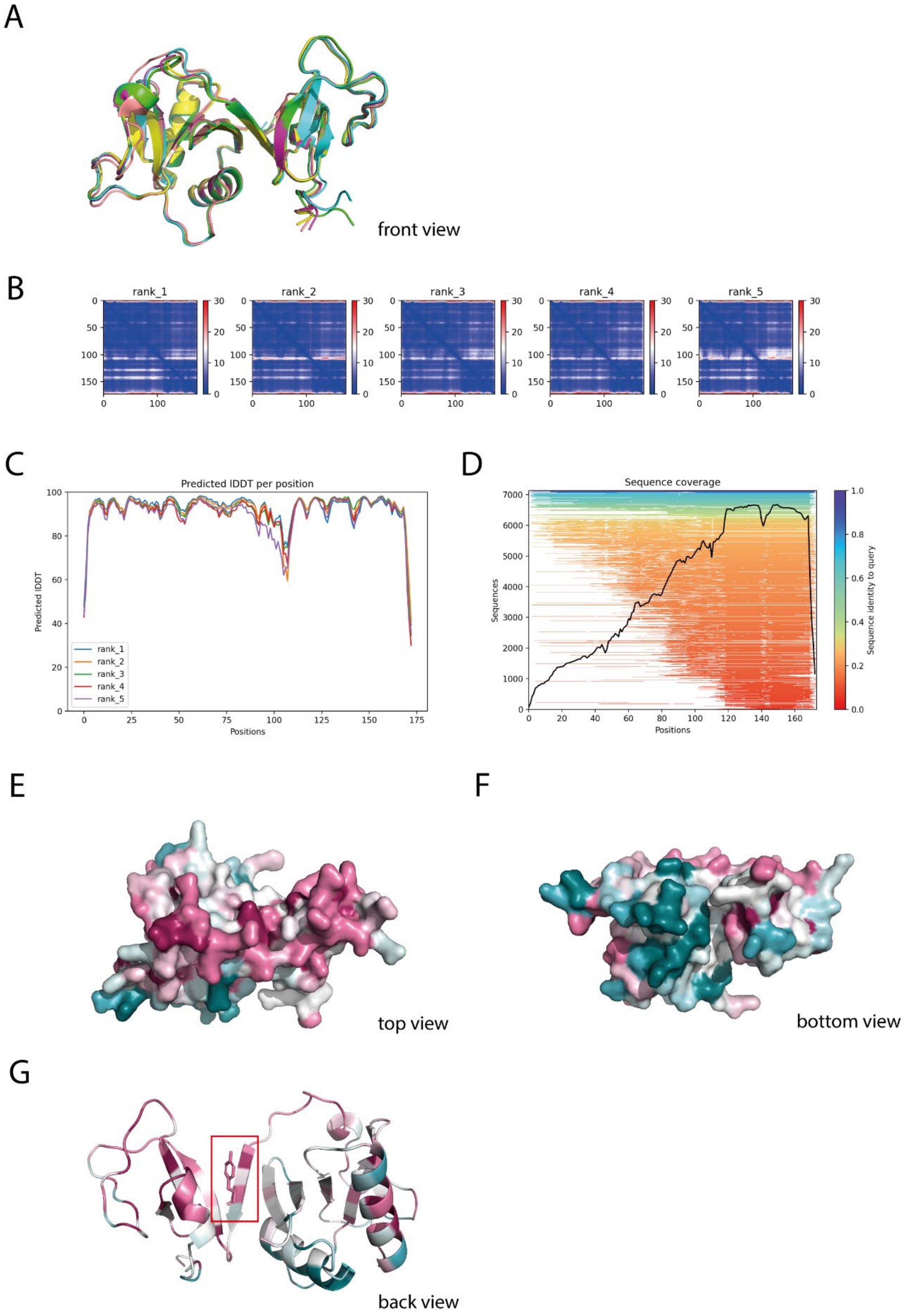

Fig S1 A. Overlay of the five top scoring Alphafold Colab models; Rank1 (green), Rank 2 (cyan), Rank 3 (purple), Rank 4 (yellow), Rank 5 (salmon). B. predicted alignment error (PAE) for the different models. C. plDDT per residue for the different models. D. sequence coverage of the multiple sequence alignment. E. Top view of the regulatory module coloured by conservation as analysed by ConSurf (Red: high conservation – Green low conservation). F. Bottom view of of the aPKCι regulatory module coloured by conservation as analyzed by ConSurf. G. cartoon representation of the aPKCι regulatory module indicating the conserved Tyr residue central to the BSL.

#### Figure S2

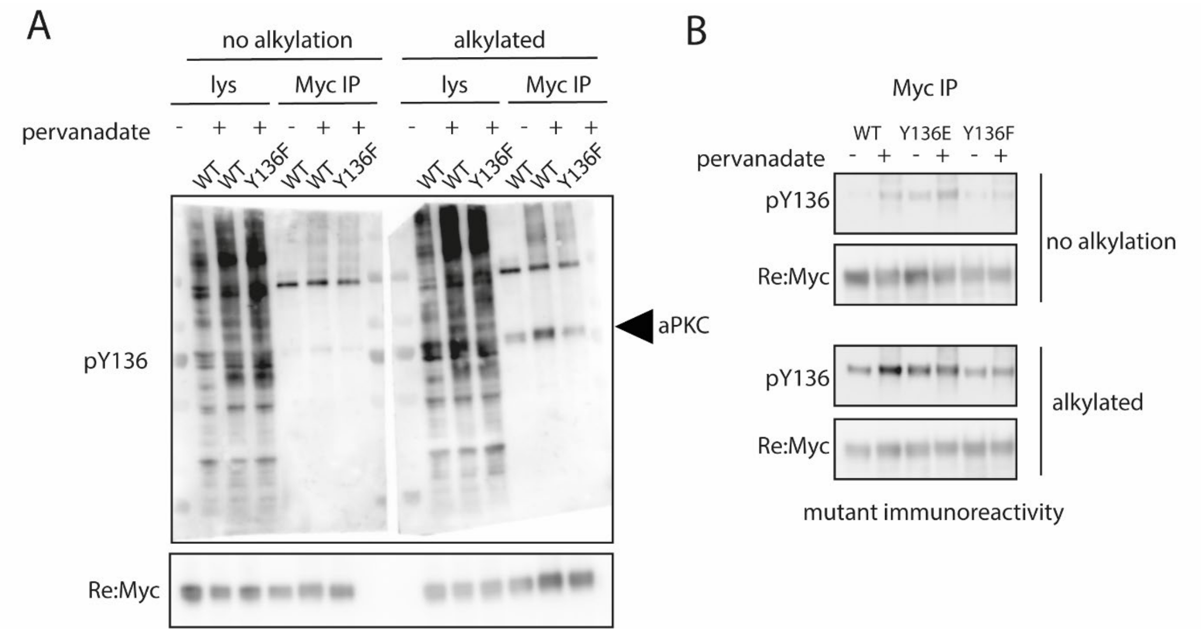

Fig S2. Specificity of the phosho-Tyr136 antibody. A. cells transfected with the indicated constructs were treated with pervanadate or left untreated. Lysates and Myc immunoprecipitates were loaded on an SDS-PAGE gel, transferred to nitrocellulose and subjected to a reduction-alkylation step or left untreated; and subsequently analysed with the pTyr-136 antibody. The antibody shows low specificity in cell lysates with broad recognition of pervanadate responsive signals, but the immunoprecipitated kinase is specifically recognized after pervanadate stimulation only when Tyr-136 is intact and post-alkylation. B. Validation of the specificity of the antibody towards immunoprecipitated aPKC and mutants with or without prior alkylation.

Figure S3

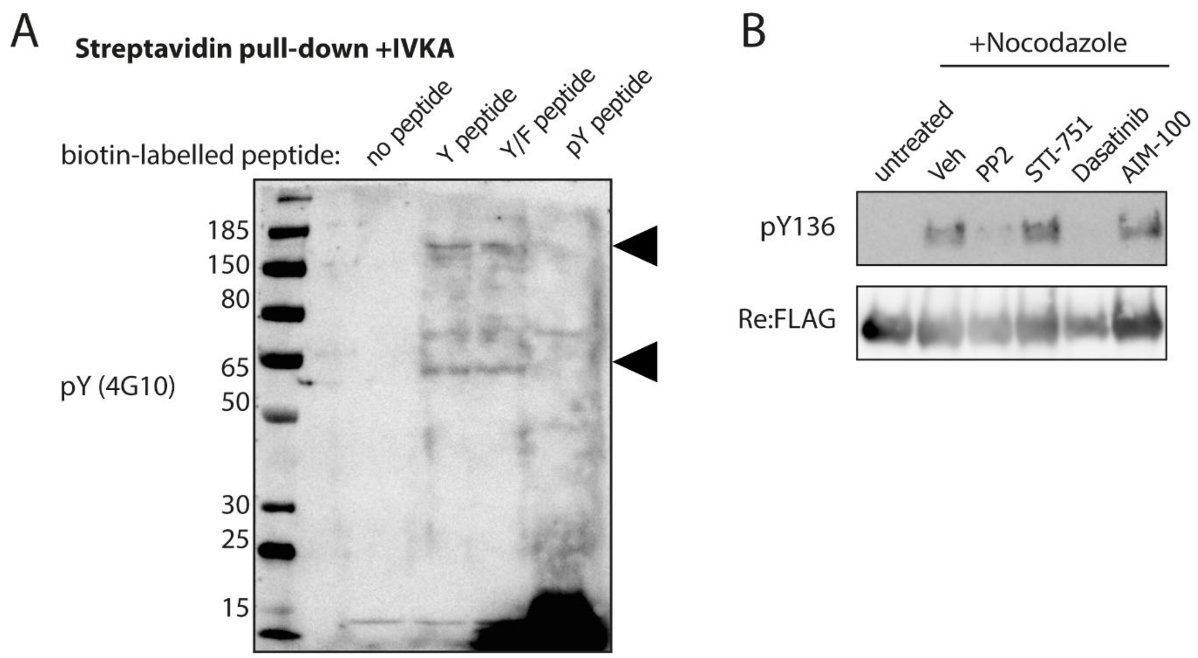

Fig S3. A. Tyr phosphorylation activity detection after precipitation of proteins from cell extract with the indicated pseudo-substrate peptides. The precipitates were spiked with ATP.$Mg^{2+}$ for 60' and tyrosine (auto)phosphorylation was followed with the pY (4G10) antibody. Bands were detected for non-phosphorylated an Y/F peptide around ~60 and ~170 kDa (arrowheads). B. Responsiveness of the Tyr-136 phospho-signal to Tyr kinase inhibitors in the context of the $aPKC\iota^{RM}$. Flag-tagged regulatory region was expressed and left untreated or treated with nocodazole and the indicated compounds. The $aPKC\iota^{RM}$ was isolated via FLAG IP and pTyr-136 levels were analyzed via Western blot.

### Table S1

#### MODA per-residue prediction scores

| num | act no | name | sel | nonStandard | counted | plainMODA | curvIndex | curvMODA | sPlainMODA | sCurvMODA |
|---|---|---|---|---|---|---|---|---|---|---|
| 1 | 14 | val | AF_aPKCi_RegDom.a/^V14 | 0 | 1 | 0 | 0.955511 | 0.00 | 2.68E-09 | 2.56E-09 |
| 2 | 15 | ala | AF_aPKCi_RegDom.a/^A15 | 0 | 1 | 0 | 0.955511 | 0.00 | 1.39E-07 | 1.33E-07 |
| 3 | 16 | gly | AF_aPKCi_RegDom.a/^G16 | 0 | 1 | 0 | 0.955511 | 0.00 | 7.39E-06 | 7.07E-06 |
| 4 | 17 | gly | AF_aPKCi_RegDom.a/^G17 | 0 | 1 | 0 | 0.955511 | 0.00 | 3.38E-04 | 3.23E-04 |
| 5 | 18 | gly | AF_aPKCi_RegDom.a/^G18 | 0 | 1 | 0 | 0.955511 | 0.00 | 3.08E-03 | 2.94E-03 |
| 6 | 19 | ser | AF_aPKCi_RegDom.a/^S19 | 0 | 1 | 0 | 0.955511 | 0.00 | 1.17E-02 | 1.12E-02 |
| 7 | 20 | gly | AF_aPKCi_RegDom.a/^G20 | 0 | 1 | 0 | 0.955511 | 0.00 | 1.43E-02 | 1.36E-02 |
| 8 | 21 | asp | AF_aPKCi_RegDom.a/^D21 | 0 | 1 | 0 | 0.955511 | 0.00 | 2.82E-03 | 2.69E-03 |
| 9 | 22 | his | AF_aPKCi_RegDom.a/^H22 | 0 | 1 | 0 | 0.863908 | 0.00 | 2.24E-01 | 1.93E-01 |
| 10 | 23 | ser | AF_aPKCi_RegDom.a/^S23 | 0 | 1 | 0 | 0.955511 | 0.00 | 1.60E-02 | 1.53E-02 |
| 11 | 24 | his | AF_aPKCi_RegDom.a/^H24 | 0 | 1 | 0 | 0.771423 | 0.00 | 1.79E-03 | 1.38E-03 |
| 12 | 25 | gln | AF_aPKCi_RegDom.a/^Q25 | 0 | 1 | 0 | 0.442339 | 0.00 | 1.09E-01 | 4.80E-02 |
| 13 | 26 | val | AF_aPKCi_RegDom.a/^V26 | 0 | 0 | 0 | 0.22995 | 0.00 | 3.84E-02 | 8.84E-03 |
| 14 | 27 | arg | AF_aPKCi_RegDom.a/^R27 | 0 | 1 | 0 | 0.121592 | 0.00 | 7.10E-01 | 8.63E-02 |
| 15 | 28 | val | AF_aPKCi_RegDom.a/^V28 | 0 | 0 | 0 | -0.06125 | 0.00 | 2.94E-01 | -1.80E-02 |
| 16 | 29 | lys | AF_aPKCi_RegDom.a/^K29 | 0 | 1 | 0 | -0.01848 | 0.00 | 3.70E+00 | -6.84E-02 |
| 17 | 30 | ala | AF_aPKCi_RegDom.a/^A30 | 0 | 0 | 0 | -0.12028 | 0.00 | 2.09E+00 | -2.51E-01 |
| 18 | 31 | tyr | AF_aPKCi_RegDom.a/^Y31 | 0 | 0 | 0 | 0.078894 | 0.00 | 9.35E+00 | 7.38E-01 |
| 19 | 32 | tyr | AF_aPKCi_RegDom.a/^Y32 | 0 | 1 | 0 | 0.0048 | 0.00 | 7.96E+00 | 3.82E-02 |
| 20 | 33 | arg | AF_aPKCi_RegDom.a/^R33 | 0 | 1 | 8.75877 | 0.16108 | 1.41 | 1.72E+01 | 2.77E+00 |
| 21 | 34 | gly | AF_aPKCi_RegDom.a/^G34 | 0 | 1 | 50.3754 | 0.250359 | 12.61 | 3.81E+01 | 9.54E+00 |
| 22 | 35 | asp | AF_aPKCi_RegDom.a/^D35 | 0 | 1 | 69.2394 | 0.056991 | 3.95 | 2.57E+01 | 1.46E+00 |
| 23 | 36 | ile | AF_aPKCi_RegDom.a/^I36 | 0 | 0 | 0 | 0.046608 | 0.00 | 2.08E+01 | 9.67E-01 |
| 24 | 37 | met | AF_aPKCi_RegDom.a/^M37 | 0 | 0 | 0 | -0.0981 | 0.00 | 6.18E+00 | -6.06E-01 |
| 25 | 38 | ile | AF_aPKCi_RegDom.a/^I38 | 0 | 0 | 0 | -0.08037 | 0.00 | 9.45E+00 | -7.59E-01 |
| 26 | 39 | thr | AF_aPKCi_RegDom.a/^T39 | 0 | 0 | 0 | -0.06291 | 0.00 | 1.27E+00 | -7.99E-02 |
| 27 | 40 | his | AF_aPKCi_RegDom.a/^H40 | 0 | 1 | 0 | 0.169387 | 0.00 | 8.07E-01 | 1.37E-01 |
| 28 | 41 | phe | AF_aPKCi_RegDom.a/^F41 | 0 | 0 | 0 | 0.1265 | 0.00 | 1.67E-01 | 2.11E-02 |
| 29 | 42 | glu | AF_aPKCi_RegDom.a/^E42 | 0 | 1 | 0 | 0.456775 | 0.00 | 5.49E-02 | 2.51E-02 |
| 30 | 43 | pro | AF_aPKCi_RegDom.a/^P43 | 0 | 1 | 0 | 0.413076 | 0.00 | 7.75E-02 | 3.20E-02 |
| 31 | 44 | ser | AF_aPKCi_RegDom.a/^S44 | 0 | 1 | 0 | 0.456887 | 0.00 | 2.53E-01 | 1.16E-01 |
| 32 | 45 | ile | AF_aPKCi_RegDom.a/^I45 | 0 | 1 | 0 | 0.182552 | 0.00 | 1.04E+00 | 1.91E-01 |
| 33 | 46 | ser | AF_aPKCi_RegDom.a/^S46 | 0 | 1 | 0 | 0.257604 | 0.00 | 3.91E+00 | 1.01E+00 |
| 34 | 47 | phe | AF_aPKCi_RegDom.a/^F47 | 0 | 1 | 34.4018 | 0.178115 | 6.13 | 9.24E+00 | 1.65E+00 |
| 35 | 48 | glu | AF_aPKCi_RegDom.a/^E48 | 0 | 1 | 0 | 0.253927 | 0.00 | 4.21E+00 | 1.07E+00 |
| 36 | 49 | gly | AF_aPKCi_RegDom.a/^G49 | 0 | 1 | 0 | 0.159268 | 0.00 | 1.81E+00 | 2.89E-01 |
| 37 | 50 | leu | AF_aPKCi_RegDom.a/^L50 | 0 | 1 | 0 | 0.02849 | 0.00 | 2.61E+00 | 7.42E-02 |
| 38 | 51 | cys | AF_aPKCi_RegDom.a/^C51 | 0 | 1 | 0 | 0.134612 | 0.00 | 2.77E+00 | 3.72E-01 |
| 39 | 52 | asn | AF_aPKCi_RegDom.a/^N52 | 0 | 1 | 0 | 0.191554 | 0.00 | 4.94E-01 | 9.47E-02 |
| 40 | 53 | glu | AF_aPKCi_RegDom.a/^E53 | 0 | 1 | 0 | 0.011378 | 0.00 | 2.87E-01 | 3.27E-03 |
| 41 | 54 | val | AF_aPKCi_RegDom.a/^V54 | 0 | 0 | 0 | -0.0316 | 0.00 | 4.98E-01 | -1.57E-02 |
| 42 | 55 | arg | AF_aPKCi_RegDom.a/^R55 | 0 | 1 | 0 | 0.206477 | 0.00 | 1.17E-01 | 2.43E-02 |
| 43 | 56 | asp | AF_aPKCi_RegDom.a/^D56 | 0 | 1 | 0 | 0.139246 | 0.00 | 1.22E-01 | 1.69E-02 |
| 44 | 57 | met | AF_aPKCi_RegDom.a/^M57 | 0 | 1 | 0 | -0.04615 | 0.00 | 1.34E+00 | -6.20E-02 |
| 45 | 58 | cys | AF_aPKCi_RegDom.a/^C58 | 0 | 1 | 0 | 0.03953 | 0.00 | 1.83E+00 | 7.25E-02 |
| 46 | 59 | ser | AF_aPKCi_RegDom.a/^S59 | 0 | 1 | 0 | 0.219373 | 0.00 | 1.07E+00 | 2.35E-01 |
| 47 | 60 | phe | AF_aPKCi_RegDom.a/^F60 | 0 | 1 | 0 | 0.26034 | 0.00 | 1.49E-01 | 3.88E-02 |
| 48 | 61 | asp | AF_aPKCi_RegDom.a/^D61 | 0 | 1 | 0 | 0.574325 | 0.00 | 8.34E-03 | 4.79E-03 |
| 49 | 62 | asn | AF_aPKCi_RegDom.a/^N62 | 0 | 1 | 0 | 0.600527 | 0.00 | 1.34E-02 | 8.03E-03 |
| 50 | 63 | glu | AF_aPKCi_RegDom.a/^E63 | 0 | 1 | 0 | 0.699709 | 0.00 | 8.09E-03 | 5.66E-03 |
| 51 | 64 | gln | AF_aPKCi_RegDom.a/^Q64 | 0 | 1 | 0 | 0.430638 | 0.00 | 3.97E-02 | 1.71E-02 |
| 52 | 65 | leu | AF_aPKCi_RegDom.a/^L65 | 0 | 1 | 0 | 0.407663 | 0.00 | 4.13E-01 | 1.69E-01 |
| 53 | 66 | phe | AF_aPKCi_RegDom.a/^F66 | 0 | 1 | 0 | 0.155678 | 0.00 | 4.76E-01 | 7.41E-02 |
| 54 | 67 | thr | AF_aPKCi_RegDom.a/^T67 | 0 | 0 | 0 | 0.204991 | 0.00 | 1.48E+00 | 3.04E-01 |
| 55 | 68 | met | AF_aPKCi_RegDom.a/^M68 | 0 | 0 | 0 | -0.00397 | 0.00 | 2.32E+00 | -9.19E-03 |
| 56 | 69 | lys | AF_aPKCi_RegDom.a/^K69 | 0 | 0 | 0 | 0.144939 | 0.00 | 8.63E-01 | 1.25E-01 |
| 57 | 70 | trp | AF_aPKCi_RegDom.a/^W70 | 0 | 1 | 0 | 0.09659 | 0.00 | 9.11E-01 | 8.80E-02 |
| 58 | 71 | ile | AF_aPKCi_RegDom.a/^I71 | 0 | 1 | 0 | 0.330405 | 0.00 | 2.83E+00 | 9.35E-01 |
| 59 | 72 | asp | AF_aPKCi_RegDom.a/^D72 | 0 | 1 | 0 | 0.433226 | 0.00 | 3.58E-01 | 1.55E-01 |
| 60 | 73 | glu | AF_aPKCi_RegDom.a/^E73 | 0 | 1 | 0 | 0.620579 | 0.00 | 1.74E-01 | 1.08E-01 |
| 61 | 74 | glu | AF_aPKCi_RegDom.a/^E74 | 0 | 1 | 0 | 0.675797 | 0.00 | 1.88E-01 | 1.27E-01 |
| 62 | 75 | gly | AF_aPKCi_RegDom.a/^G75 | 0 | 1 | 0 | 0.576345 | 0.00 | 2.97E+00 | 1.71E+00 |
| 63 | 76 | asp | AF_aPKCi_RegDom.a/^D76 | 0 | 1 | 0 | 0.531604 | 0.00 | 3.38E-01 | 1.80E-01 |
| 64 | 77 | pro | AF_aPKCi_RegDom.a/^P77 | 0 | 1 | 0 | 0.335446 | 0.00 | 6.62E-01 | 2.22E-01 |
| 65 | 78 | cys | AF_aPKCi_RegDom.a/^C78 | 0 | 1 | 0 | 0.284198 | 0.00 | 1.87E+00 | 5.31E-01 |
| 66 | 79 | thr | AF_aPKCi_RegDom.a/^T79 | 0 | 1 | 0 | 0.259678 | 0.00 | 5.91E+00 | 1.54E+00 |
| 67 | 80 | val | AF_aPKCi_RegDom.a/^V80 | 0 | 0 | 0 | 0.065274 | 0.00 | 6.12E+00 | 4.00E-01 |
| 68 | 81 | ser | AF_aPKCi_RegDom.a/^S81 | 0 | 1 | 46.1953 | 0.264052 | 12.20 | 1.65E+01 | 4.36E+00 |
| 69 | 82 | ser | AF_aPKCi_RegDom.a/^S82 | 0 | 1 | 24.2305 | 0.293989 | 7.12 | 1.59E+01 | 4.67E+00 |
| 70 | 83 | gln | AF_aPKCi_RegDom.a/^Q83 | 0 | 1 | 0 | 0.290936 | 0.00 | 8.68E+00 | 2.53E+00 |
| 71 | 84 | leu | AF_aPKCi_RegDom.a/^L84 | 0 | 1 | 0 | 0.348949 | 0.00 | 8.19E+00 | 2.86E+00 |
| 72 | 85 | glu | AF_aPKCi_RegDom.a/^E85 | 0 | 1 | 23.3118 | 0.222764 | 5.19 | 1.01E+01 | 2.24E+00 |
| 73 | 86 | leu | AF_aPKCi_RegDom.a/^L86 | 0 | 0 | 0 | 0.140063 | 0.00 | 5.92E+00 | 8.29E-01 |
| 74 | 87 | glu | AF_aPKCi_RegDom.a/^E87 | 0 | 1 | 0 | 0.302841 | 0.00 | 2.76E+00 | 8.34E-01 |
| 75 | 88 | glu | AF_aPKCi_RegDom.a/^E88 | 0 | 1 | 0 | 0.316468 | 0.00 | 2.81E+00 | 8.89E-01 |
| 76 | 89 | ala | AF_aPKCi_RegDom.a/^A89 | 0 | 0 | 0 | 0.158305 | 0.00 | 1.71E+00 | 2.71E-01 |
| 77 | 90 | phe | AF_aPKCi_RegDom.a/^F90 | 0 | 1 | 0 | 0.259471 | 0.00 | 5.17E-01 | 1.34E-01 |
| 78 | 91 | arg | AF_aPKCi_RegDom.a/^R91 | 0 | 1 | 0 | 0.411495 | 0.00 | 3.07E-01 | 1.26E-01 |
| 79 | 92 | leu | AF_aPKCi_RegDom.a/^L92 | 0 | 1 | 0 | 0.416168 | 0.00 | 2.00E-01 | 8.34E-02 |
| 80 | 93 | tyr | AF_aPKCi_RegDom.a/^Y93 | 0 | 1 | 0 | 0.310967 | 0.00 | 7.18E-02 | 2.23E-02 |
| 81 | 94 | glu | AF_aPKCi_RegDom.a/^E94 | 0 | 1 | 0 | 0.509005 | 0.00 | 1.18E-02 | 6.02E-03 |
| 82 | 95 | leu | AF_aPKCi_RegDom.a/^L95 | 0 | 1 | 0 | 0.602356 | 0.00 | 1.00E-02 | 6.05E-03 |
| 83 | 96 | asn | AF_aPKCi_RegDom.a/^N96 | 0 | 1 | 0 | 0.584492 | 0.00 | 8.30E-02 | 4.85E-02 |
| 84 | 97 | lys | AF_aPKCi_RegDom.a/^K97 | 0 | 1 | 0 | 0.68372 | 0.00 | 1.15E-02 | 7.87E-03 |
| 85 | 98 | asp | AF_aPKCi_RegDom.a/^D98 | 0 | 1 | 0 | 0.411447 | 0.00 | 3.03E-01 | 1.25E-01 |
| 86 | 99 | ser | AF_aPKCi_RegDom.a/^S99 | 0 | 1 | 0 | 0.400728 | 0.00 | 4.54E-01 | 1.82E-01 |
| 87 | 100 | glu | AF_aPKCi_RegDom.a/^E100 | 0 | 1 | 0 | 0.14435 | 0.00 | 7.48E-01 | 1.08E-01 |
| 88 | 101 | leu | AF_aPKCi_RegDom.a/^L101 | 0 | 0 | 0 | 0.073632 | 0.00 | 1.62E-01 | 1.20E-02 |
| 89 | 102 | leu | AF_aPKCi_RegDom.a/^L102 | 0 | 0 | 0 | 0.124071 | 0.00 | 3.19E+00 | 3.95E-01 |
| 90 | 103 | ile | AF_aPKCi_RegDom.a/^I103 | 0 | 0 | 0 | -0.0747 | 0.00 | 5.89E-01 | -4.40E-02 |
| 91 | 104 | his | AF_aPKCi_RegDom.a/^H104 | 0 | 0 | 0 | 0.094057 | 0.00 | 1.61E+00 | 1.51E-01 |
| 92 | 105 | val | AF_aPKCi_RegDom.a/^V105 | 0 | 0 | 0 | -0.02363 | 0.00 | 1.42E+00 | -3.35E-02 |
| 93 | 106 | phe | AF_aPKCi_RegDom.a/^F106 | 0 | 0 | 0 | 0.189712 | 0.00 | 4.63E-01 | 8.78E-02 |
| 94 | 107 | pro | AF_aPKCi_RegDom.a/^P107 | 0 | 1 | 0 | 0.221431 | 0.00 | 8.51E-01 | 1.89E-01 |
| 95 | 108 | cys | AF_aPKCi_RegDom.a/^C108 | 0 | 1 | 0 | 0.367745 | 0.00 | 1.13E-01 | 4.14E-02 |
| 96 | 109 | val | AF_aPKCi_RegDom.a/^V109 | 0 | 1 | 0 | 0.442476 | 0.00 | 9.93E-01 | 4.39E-01 |
| 97 | 110 | pro | AF_aPKCi_RegDom.a/^P110 | 0 | 1 | 0 | 0.539266 | 0.00 | 6.90E-01 | 3.72E-01 |
| 98 | 111 | glu | AF_aPKCi_RegDom.a/^E111 | 0 | 1 | 0 | 0.764629 | 0.00 | 2.17E-01 | 1.66E-01 |
| 99 | 112 | arg | AF_aPKCi_RegDom.a/^R112 | 0 | 1 | 0 | 0.846549 | 0.00 | 1.00E+00 | 8.48E-01 |
| 100 | 113 | pro | AF_aPKCi_RegDom.a/^P113 | 0 | 1 | 6.67797 | 0.71527 | 4.78 | 3.99E+00 | 2.86E+00 |
| 101 | 114 | gly | AF_aPKCi_RegDom.a/^G114 | 0 | 1 | 0 | 0.64435 | 0.00 | 1.53E+00 | 9.84E-01 |
| 102 | 115 | met | AF_aPKCi_RegDom.a/^M115 | 0 | 1 | 0 | 0.695384 | 0.00 | 4.11E-01 | 2.86E-01 |
| 103 | 116 | pro | AF_aPKCi_RegDom.a/^P116 | 0 | 1 | 0 | 0.533538 | 0.00 | 9.65E-02 | 5.15E-02 |
| 104 | 117 | cys | AF_aPKCi_RegDom.a/^C117 | 0 | 1 | 0 | 0.365372 | 0.00 | 1.64E-01 | 5.98E-02 |
| 105 | 118 | pro | AF_aPKCi_RegDom.a/^P118 | 0 | 1 | 0 | 0.521566 | 0.00 | 3.41E-02 | 1.78E-02 |
| 106 | 119 | gly | AF_aPKCi_RegDom.a/^G119 | 0 | 1 | 0 | 0.481899 | 0.00 | 3.04E-01 | 1.47E-01 |
| 107 | 120 | glu | AF_aPKCi_RegDom.a/^E120 | 0 | 1 | 0 | 0.407149 | 0.00 | 2.77E-01 | 1.13E-01 |

| | | | | | | | | | | | |
|---|---|---|---|---|---|---|---|---|---|---|---|
| 108 | 121 | asp | AF_aPKCi_RegDom.a/^D121 | 0 | 1 | 0 | 0.564555 | 0.00 | 2.73E+00 | 1.54E+00 | |
| 109 | 122 | lys | AF_aPKCi_RegDom.a/^K122 | 0 | 1 | 0 | 0.592906 | 0.00 | 3.74E+00 | 2.22E+00 | |
| 110 | 123 | ser | AF_aPKCi_RegDom.a/^S123 | 0 | 1 | 0 | 0.690659 | 0.00 | 1.44E+01 | 9.92E+00 | |
| 111 | 124 | ile | AF_aPKCi_RegDom.a/^I124 | 0 | 1 | 0 | 0.444652 | 0.00 | 1.20E+01 | 5.35E+00 | |
| 112 | 125 | tyr | AF_aPKCi_RegDom.a/^Y125 | 0 | 1 | 0 | 0.462707 | 0.00 | 1.15E+01 | 5.34E+00 | MUTANTS |
| 113 | 126 | arg | AF_aPKCi_RegDom.a/^R126 | 0 | 1 | 62.0191 | 0.735017 | 45.59 | 4.95E+01 | 3.64E+01 | Ala |
| 114 | 127 | arg | AF_aPKCi_RegDom.a/^R127 | 0 | 1 | 128.266 | 0.70799 | 90.81 | 9.64E+01 | 6.83E+01 | Ala |
| 115 | 128 | gly | AF_aPKCi_RegDom.a/^G128 | 0 | 1 | 190.937 | 0.787455 | 150.35 | 1.18E+02 | 9.29E+01 | |
| 116 | 129 | ala | AF_aPKCi_RegDom.a/^A129 | 0 | 1 | 25.8598 | 0.742808 | 19.21 | 5.60E+01 | 4.16E+01 | |
| 117 | 130 | arg | AF_aPKCi_RegDom.a/^R130 | 0 | 1 | 15.5355 | 0.586375 | 9.11 | 7.13E+01 | 4.18E+01 | Ala |
| 118 | 131 | arg | AF_aPKCi_RegDom.a/^R131 | 0 | 1 | 400.232 | 0.607438 | 243.12 | 1.77E+02 | 1.08E+02 | Ala |
| 119 | 132 | trp | AF_aPKCi_RegDom.a/^W132 | 0 | 1 | 37.262 | 0.284438 | 10.60 | 5.75E+01 | 1.64E+01 | |
| 120 | 133 | arg | AF_aPKCi_RegDom.a/^R133 | 0 | 1 | 30.7195 | 0.181657 | 5.58 | 4.81E+01 | 8.74E+00 | |
| 121 | 134 | lys | AF_aPKCi_RegDom.a/^K134 | 0 | 1 | 40.0413 | 0.077114 | 3.09 | 4.10E+01 | 3.16E+00 | |
| 122 | 135 | leu | AF_aPKCi_RegDom.a/^L135 | 0 | 0 | 0 | -0.02991 | 0.00 | 1.66E+01 | -4.97E-01 | |
| 123 | 136 | tyr | AF_aPKCi_RegDom.a/^Y136 | 0 | 1 | 14.6314 | -0.03825 | -0.56 | 3.05E+01 | -1.17E+00 | |
| 124 | 137 | cys | AF_aPKCi_RegDom.a/^C137 | 0 | 0 | 0 | -0.04303 | 0.00 | 3.55E+00 | -1.53E-01 | |
| 125 | 138 | ala | AF_aPKCi_RegDom.a/^A138 | 0 | 0 | 0 | 0.112779 | 0.00 | 7.20E+00 | 8.12E-01 | |
| 126 | 139 | asn | AF_aPKCi_RegDom.a/^N139 | 0 | 1 | 0 | 0.298443 | 0.00 | 1.21E+00 | 3.60E-01 | |
| 127 | 140 | gly | AF_aPKCi_RegDom.a/^G140 | 0 | 1 | 0 | 0.220739 | 0.00 | 8.94E-01 | 1.97E-01 | |
| 128 | 141 | his | AF_aPKCi_RegDom.a/^H141 | 0 | 1 | 0 | 0.156672 | 0.00 | 6.88E+00 | 1.08E+00 | |
| 129 | 142 | thr | AF_aPKCi_RegDom.a/^T142 | 0 | 0 | 0 | 0.047584 | 0.00 | 6.35E+00 | 3.02E-01 | |
| 130 | 143 | phe | AF_aPKCi_RegDom.a/^F143 | 0 | 0 | 0 | 0.029121 | 0.00 | 3.54E+01 | 1.03E+00 | MUTANTS |
| 131 | 144 | gln | AF_aPKCi_RegDom.a/^Q144 | 0 | 1 | 9.59911 | 0.09059 | 0.87 | 4.30E+01 | 3.89E+00 | |
| 132 | 145 | ala | AF_aPKCi_RegDom.a/^A145 | 0 | 1 | 79.648 | 0.170145 | 13.55 | 1.50E+02 | 2.55E+01 | |
| 133 | 146 | lys | AF_aPKCi_RegDom.a/^K146 | 0 | 1 | 5.28948 | 0.296912 | 1.57 | 1.46E+02 | 4.34E+01 | |
| 134 | 147 | arg | AF_aPKCi_RegDom.a/^R147 | 0 | 1 | 525.681 | 0.568803 | 299.01 | 4.03E+02 | 2.29E+02 | Ala |
| 135 | 148 | phe | AF_aPKCi_RegDom.a/^F148 | 0 | 1 | 129.572 | 0.512654 | 66.43 | 2.34E+02 | 1.20E+02 | |
| 136 | 149 | asn | AF_aPKCi_RegDom.a/^N149 | 0 | 1 | 252.267 | 0.76876 | 193.93 | 2.84E+02 | 2.18E+02 | |
| 137 | 150 | arg | AF_aPKCi_RegDom.a/^R150 | 0 | 1 | 190.857 | 0.732672 | 139.84 | 2.94E+02 | 2.15E+02 | Ala |
| 138 | 151 | arg | AF_aPKCi_RegDom.a/^R151 | 0 | 1 | 882.099 | 0.727214 | 641.48 | 4.62E+02 | 3.36E+02 | Ala |
| 139 | 152 | ala | AF_aPKCi_RegDom.a/^A152 | 0 | 1 | 0 | 0.477964 | 0.00 | 1.93E+02 | 9.22E+01 | |
| 140 | 153 | his | AF_aPKCi_RegDom.a/^H153 | 0 | 1 | 117.466 | 0.485148 | 56.99 | 1.57E+02 | 7.62E+01 | |
| 141 | 154 | cys | AF_aPKCi_RegDom.a/^C154 | 0 | 1 | 0 | 0.329839 | 0.00 | 5.15E+01 | 1.70E+01 | |
| 142 | 155 | ala | AF_aPKCi_RegDom.a/^A155 | 0 | 1 | 0.722752 | 0.308711 | 0.22 | 1.70E+01 | 5.25E+00 | |
| 143 | 156 | ile | AF_aPKCi_RegDom.a/^I156 | 0 | 1 | 2.80178 | 0.3392 | 0.95 | 8.87E+00 | 3.01E+00 | |
| 144 | 157 | cys | AF_aPKCi_RegDom.a/^C157 | 0 | 1 | 0 | 0.429731 | 0.00 | 2.01E+01 | 8.65E+00 | |
| 145 | 158 | thr | AF_aPKCi_RegDom.a/^T158 | 0 | 1 | 0 | 0.516691 | 0.00 | 4.40E+01 | 2.27E+01 | |
| 146 | 159 | asp | AF_aPKCi_RegDom.a/^D159 | 0 | 1 | 0 | 0.507618 | 0.00 | 1.31E+02 | 6.64E+01 | MUTANTS |
| 147 | 160 | arg | AF_aPKCi_RegDom.a/^R160 | 0 | 1 | 632.666 | 0.602369 | 381.10 | 3.92E+02 | 2.36E+02 | Ala |
| 148 | 161 | ile | AF_aPKCi_RegDom.a/^I161 | 0 | 0 | 0 | 0.412062 | 0.00 | 2.07E+02 | 8.52E+01 | |
| 149 | 162 | trp | AF_aPKCi_RegDom.a/^W162 | 0 | 1 | 1238.04 | 0.646716 | 800.66 | 6.05E+02 | 3.91E+02 | Ala |
| 150 | 163 | gly | AF_aPKCi_RegDom.a/^G163 | 0 | 1 | 772.348 | 0.679519 | 524.83 | 7.31E+02 | 4.96E+02 | |
| 151 | 164 | leu | AF_aPKCi_RegDom.a/^L164 | 0 | 1 | 1233.21 | 0.822926 | 1014.84 | 9.40E+02 | 7.73E+02 | Asp |
| 152 | 165 | gly | AF_aPKCi_RegDom.a/^G165 | 0 | 1 | 997.332 | 0.656699 | 654.95 | 7.00E+02 | 4.60E+02 | |
| 153 | 166 | arg | AF_aPKCi_RegDom.a/^R166 | 0 | 1 | 435.699 | 0.385039 | 167.76 | 3.51E+02 | 1.35E+02 | |
| 154 | 167 | gln | AF_aPKCi_RegDom.a/^Q167 | 0 | 1 | 396.592 | 0.480482 | 190.56 | 3.86E+02 | 1.85E+02 | |
| 155 | 168 | gly | AF_aPKCi_RegDom.a/^G168 | 0 | 0 | 0 | 0.253542 | 0.00 | 1.57E+02 | 3.98E+01 | |
| 156 | 169 | tyr | AF_aPKCi_RegDom.a/^Y169 | 0 | 1 | 271.095 | 0.091134 | 24.71 | 7.30E+01 | 6.65E+00 | |
| 157 | 170 | lys | AF_aPKCi_RegDom.a/^K170 | 0 | 1 | 12.6393 | 0.185104 | 2.34 | 4.97E+01 | 9.20E+00 | |
| 158 | 171 | cys | AF_aPKCi_RegDom.a/^C171 | 0 | 0 | 0 | 0.160665 | 0.00 | 1.38E+01 | 2.22E+00 | |
| 159 | 172 | ile | AF_aPKCi_RegDom.a/^I172 | 0 | 1 | 0 | 0.125817 | 0.00 | 9.89E+00 | 1.24E+00 | |
| 160 | 173 | asn | AF_aPKCi_RegDom.a/^N173 | 0 | 1 | 0 | 0.249556 | 0.00 | 3.64E+00 | 9.09E-01 | |
| 161 | 174 | cys | AF_aPKCi_RegDom.a/^C174 | 0 | 1 | 9.67618 | 0.326067 | 3.16 | 6.74E+00 | 2.20E+00 | |
| 162 | 175 | lys | AF_aPKCi_RegDom.a/^K175 | 0 | 1 | 19.1006 | 0.288026 | 5.50 | 1.68E+01 | 4.83E+00 | |
| 163 | 176 | leu | AF_aPKCi_RegDom.a/^L176 | 0 | 1 | 10.2335 | 0.234119 | 2.40 | 1.44E+01 | 3.38E+00 | |
| 164 | 177 | leu | AF_aPKCi_RegDom.a/^L177 | 0 | 1 | 40.8581 | 0.270241 | 11.04 | 5.37E+01 | 1.45E+01 | |
| 165 | 178 | val | AF_aPKCi_RegDom.a/^V178 | 0 | 0 | 0 | 0.180225 | 0.00 | 3.35E+01 | 6.04E+00 | |
| 166 | 179 | his | AF_aPKCi_RegDom.a/^H179 | 0 | 1 | 0 | 0.274765 | 0.00 | 7.68E+01 | 2.11E+01 | |
| 167 | 180 | lys | AF_aPKCi_RegDom.a/^K180 | 0 | 1 | 276.303 | 0.185801 | 51.34 | 1.32E+02 | 2.46E+01 | |
| 168 | 181 | lys | AF_aPKCi_RegDom.a/^K181 | 0 | 1 | 16.9483 | 0.298168 | 5.05 | 5.34E+01 | 1.59E+01 | |
| 169 | 182 | cys | AF_aPKCi_RegDom.a/^C182 | 0 | 1 | 0 | 0.261992 | 0.00 | 2.30E+01 | 6.04E+00 | |
| 170 | 183 | his | AF_aPKCi_RegDom.a/^H183 | 0 | 0 | 0 | 0.096293 | 0.00 | 3.55E+01 | 3.42E+00 | |
| 171 | 184 | lys | AF_aPKCi_RegDom.a/^K184 | 0 | 1 | 0 | 0.286202 | 0.00 | 9.49E+00 | 2.72E+00 | |
| 172 | 185 | leu | AF_aPKCi_RegDom.a/^L185 | 0 | 1 | 1.49872 | 0.421593 | 0.63 | 4.90E+00 | 2.07E+00 | |
| 173 | 186 | val | AF_aPKCi_RegDom.a/^V186 | 0 | 1 | 0 | 0.261255 | 0.00 | 6.14E+00 | 1.60E+00 | |
| 174 | 187 | thr | AF_aPKCi_RegDom.a/^T187 | 0 | 1 | 2.03261 | 0.558332 | 1.13 | 1.35E+00 | 7.56E-01 | |
| 175 | 188 | ile | AF_aPKCi_RegDom.a/^I188 | 0 | 1 | 0.924768 | 0.468308 | 0.43 | 1.96E+00 | 9.17E-01 | |
| 176 | 189 | glu | AF_aPKCi_RegDom.a/^E189 | 0 | 1 | 0 | 0.493331 | 0.00 | 7.16E-01 | 3.53E-01 | |
| 177 | 190 | cys | AF_aPKCi_RegDom.a/^C190 | 0 | 1 | 0 | 0.313769 | 0.00 | 2.25E+00 | 7.06E-01 | |

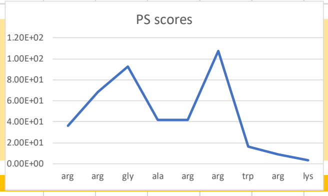

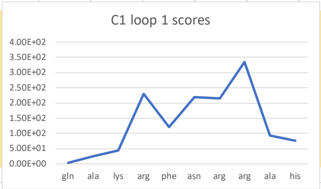

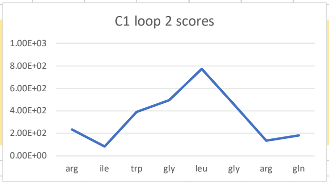

### Table S2

Amino Acid Conservation Scores
=======================================

The layers for assigning grades are as follows.
from -1.227 to -0.954 the grade is 9
from -0.954 to -0.682 the grade is 8
from -0.682 to -0.409 the grade is 7
from -0.409 to -0.136 the grade is 6
from -0.136 to 0.290 the grade is 5
from 0.290 to 0.869 the grade is 4
from 0.869 to 1.449 the grade is 3
from 1.449 to 2.028 the grade is 2
from 2.028 to 2.608 the grade is 1

If the difference between the colors of the CONFIDENCE INTERVAL COLORS is more than 3 or the msa number (under the column titled MSA) is less than 6, there is insufficient data and an * appears i

#NAME?
#NAME?
- 3LATOM: The ATOM derived sequence in three letter code, including the AA's positions as they appear in the PDB file and the chain identifier.
#NAME?
- COLOR: The color scale representing the conservation scores (9 - conserved, 1 - variable).
#NAME?
#NAME?
- MSA DATA: The number of aligned sequences having an amino acid (non-gapped) from the overall number of sequences at each position.
#NAME?

| POS | SEQ | 3LATOM | SCORE | COLOR | CONFIDEN | CONFIDEN | B/E | FUN | MSA DATA | RESIDUE VARIETY | | | | |
|---|---|---|---|---|---|---|---|---|---|---|---|---|---|---|
| | | | (normaliz | | | | | | | | | | | |
| 1 | H | HIS:1:A | -0.559 | 7 | -0.972, -0.330 | | | | 9,6 | | e | | 12/150 | H,N |
| 2 | S | SER:2:A | 0.249 | 5 | -0.256, 0.560 | | | | 6,4 | | e | | 26/150 | S,T,K,E,L,G,N |
| 3 | H | HIS:3:A | 1.411 | 3 | 0.775, 1.497 | | | | 4,2 | | e | | 48/150 | H,E,D,N,T,S,A,G,M,K |
| 4 | Q | GLN:4:A | 2.144 | 1 | 1.060, 2.631 | | | | 3,1 | | e | | 74/150 | Q,A,V,D,E,W,S,R,Y,L,K,F,G,H |
| 5 | V | VAL:5:A | -0.238 | 6 | -0.466, -0.086 | | | | 7,5 | | b | | 86/150 | V,I,L,F,K |
| 6 | R | ARG:6:A | -0.022 | 5 | -0.330, 0.121 | | | | 6,5 | | e | | 88/150 | R,P,K,T,Q,S,I,F,L,A |
| 7 | V | VAL:7:A | 0.734 | 4 | 0.246, 1.060 | | | | 5,3 | | b | | 93/150 | V,L,A,I,M,F,N,T |
| 8 | K | LYS:8:A | -0.475 | 7 | -0.693, -0.330 | | | | 8,6 | | b | | 94/150 | K,R,A,Q,L,S,M |
| 9 | A | ALA:9:A | 0.348 | 4 | 0.012, 0.560 | | | | 5,4 | | b | | 95/150 | A,V,I,Y,T,Q,L,F |
| 10 | Y | TYR:10:A | 0.486 | 4 | 0.121, 0.775 | | | | 5,4 | | e | | 97/150 | Y,F,H,R,I,A,V,G,C,S,K,L,Q |
| 11 | Y | TYR:11:A | 1.288 | 3 | 0.775, 1.497 | | | | 4,2 | | b | | 102/150 | Y,F,H,L,V,T,A,R,N,S |
| 12 | R | ARG:12:A | 2.303 | 1 | 1.060, 2.631 | | | | 3,1 | | e | | 105/150 | R,D,C,L,N,S,K,G,A,F,Q,H,T |
| 13 | G | GLY:13:A | 1.09 | 3 | 0.560, 1.497 | | | | 4,2 | | e | | 122/150 | G,R,S,D,Q,P,E,K |
| 14 | D | ASP:14:A | 0.276 | 5 | -0.086, 0.560 | | | | 5,4 | | e | | 124/150 | D,E,I,V,Q,N,G,K,L,T,S,H,Y |
| 15 | I | ILE:15:A | 0.275 | 5 | 0.012, 0.389 | | | | 5,4 | | b | | 126/150 | I,V,A,M,K,D,S,L |
| 16 | M | MET:16:A | 0.252 | 5 | -0.086, 0.389 | | | | 5,4 | | b | | 127/150 | M,I,L,V,Q,F,K,T,Y,R |
| 17 | I | ILE:17:A | -0.443 | 7 | -0.640, -0.330 | | | | 7,6 | | b | | 129/150 | I,C,V,M,K,T,Y,P |
| 18 | T | THR:18:A | 0.114 | 5 | -0.174, 0.246 | | | | 6,5 | | b | | 131/150 | T,V,L,A,M,I,S,F,R |
| 19 | H | HIS:19:A | 0.206 | 5 | -0.086, 0.389 | | | | 5,4 | | e | | 136/150 | H,N,D,S,Y,I,V,T,W,A,R,E |
| 20 | F | PHE:20:A | 0.281 | 5 | -0.086, 0.560 | | | | 5,4 | | b | | 138/150 | F,M,L,T,V,I,S,A,N,Q,C |
| 21 | E | GLU:21:A | 2.469 | 1 | 1.497, 2.631 | | | | 2,1 | | e | | 137/150 | E,N,V,R,D,T,P,Q,A,H,S,K,L,G,F |
| 22 | P | PRO:22:A | 1.378 | 3 | 0.775, 1.497 | | | | 4,2 | | e | | 138/150 | P,R,A,N,S,Q,D,H,T,V,E,K,I,L,C,Y, |
| 23 | S | SER:23:A | 1.42 | 3 | 0.775, 1.497 | | | | 4,2 | | e | | 139/150 | S,G,T,A,E,P,I,H,N,V,D,K,Q,R,C,Y, |
| 24 | I | ILE:24:A | 0.686 | 4 | 0.246, 0.775 | | | | 5,4 | | e | | 139/150 | I,M,V,L,T,A,F,P,K,S,N |
| 25 | S | SER:25:A | 1.133 | 3 | 0.560, 1.497 | | | | 4,2 | | e | | 140/150 | S,T,H,N,P,V,A,D,L,E,R,I,K,Q |
| 26 | F | PHE:26:A | 1.724 | 2 | 1.060, 2.631 | | | | 3,1 | | e | | 140/150 | F,V,Y,L,S,M,I |
| 27 | E | GLU:27:A | 2.213 | 1 | 1.060, 2.631 | | | | 3,1 | | e | | 140/150 | E,D,V,A,C,P,M,H,L,N,S,K,R,T,G,Y |
| 28 | G | GLY:28:A | 2.564 | 1 | 1.497, 2.631 | | | | 2,1 | | e | | 140/150 | G,D,E,I,T,Q,A,K,S,X,M,N,P,L,R,V |
| 29 | L | LEU:29:A | -0.058 | 5 | -0.330, 0.121 | | | | 6,5 | | b | | 142/150 | L,F,M,E,V,I |
| 30 | C | CYS:30:A | 2.397 | 1 | 1.497, 2.631 | | | | 2,1 | | b | | 142/150 | C,S,V,R,Y,H,L,K,T,G,E,F,M |
| 31 | N | ASN:31:A | 2.568 | 1 | 1.497, 2.631 | | | | 2,1 | | e | | 142/150 | N,E,D,K,S,G,Q,V,R,Y,A,M,T,H |
| 32 | E | GLU:32:A | 0.305 | 4 | 0.012, 0.560 | | | | 5,4 | | e | | 142/150 | E,N,K,L,Q,V,D,T,S,I,A,H,Y,M |
| 33 | V | VAL:33:A | -0.316 | 6 | -0.527, -0.174 | | | | 7,6 | | b | | 142/150 | V,D,I,M,L |
| 34 | R | ARG:34:A | 0.595 | 4 | 0.246, 0.775 | | | | 5,4 | | e | | 142/150 | R,K,M,P,V,L,Q,C,H,E,W |
| 35 | D | ASP:35:A | 2.098 | 1 | 1.060, 2.631 | | | | 3,1 | | e | | 142/150 | D,E,T,S,G,V,F,A,R,N,Q,K,I,L |
| 36 | M | MET:36:A | -0.202 | 6 | -0.400, -0.086 | | | | 6,5 | | b | | 143/150 | M,L,V,A,I,F,T,Q,N |
| 37 | C | CYS:37:A | -0.687 | 8 | -0.884, -0.585 | | | | 8,7 | | b | | 143/150 | C,F,W,L,Y,V |
| 38 | S | SER:38:A | 1.115 | 3 | 0.560, 1.497 | | | | 4,2 | | e | | 144/150 | S,K,C,R,G,H,T,Q,N,E,V,Y,W |
| 39 | F | PHE:39:A | -0.177 | 6 | -0.400, 0.012 | | | | 6,5 | | b | | 144/150 | F,V,L,I,M,Q,S |
| 40 | D | ASP:40:A | 2.608 | 1 | 1.497, 2.631 | | | | 2,1 | | e | | 144/150 | D,M,T,P,R,G,S,Q,H,E,A,C,N,L,V,I |
| 41 | N | ASN:41:A | 2.576 | 1 | 1.497, 2.631 | | | | 2,1 | | e | | 144/150 | N,P,K,S,Q,G,V,R,M,E,A,I,H,D,T,Y |
| 42 | E | GLU:42:A | 2.567 | 1 | 1.497, 2.631 | | | | 2,1 | | e | | 145/150 | E,S,Q,K,N,P,D,H,G,R,A,Y,M,T,V |
| 43 | Q | GLN:43:A | 0.099 | 5 | -0.174, 0.246 | | | | 6,5 | | e | | 145/150 | Q,H,E,Y,T,M,D,S,R,K,C,A,P,N |
| 44 | L | LEU:44:A | 1.682 | 2 | 1.060, 2.631 | | | | 3,1 | | e | | 145/150 | L,E,P,Q,I,K,V,A,D,H,R,S,N,T |
| 45 | F | PHE:45:A | -0.495 | 7 | -0.693, -0.330 | | | | 8,6 | | b | | 145/150 | F,L,I,C,Y,V |
| 46 | T | THR:46:A | -1.174 | 9 | -1.275, -1.139 | | | | 9,9 | | b | s | 146/150 | T,N,R,M |
| 47 | M | MET:47:A | 0.05 | 5 | -0.256, 0.246 | | | | 6,5 | | b | | 147/150 | M,L,T,I,V,F,A |
| 48 | K | LYS:48:A | -0.826 | 8 | -0.972, -0.744 | | | | 9,8 | | b | | 146/150 | K,H,P,R,D,I,Q,C |
| 49 | W | TRP:49:A | -0.573 | 7 | -0.839, -0.400 | | | | 8,6 | | b | | 146/150 | W,F,C,Y,G |
| 50 | I | ILE:50:A | -0.355 | 6 | -0.585, -0.256 | | | | 7,6 | | b | | 146/150 | I,L,V,A,Y,M |
| 51 | D | ASP:51:A | -1.09 | 9 | -1.181, -1.056 | | | | 9,9 | | e | f | 147/150 | D,E,K,P,A |
| 52 | E | GLU:52:A | -0.221 | 6 | -0.466, -0.086 | | | | 7,5 | | e | | 148/150 | E,D,S,N,V,K,G,P |
| 53 | E | GLU:53:A | -0.606 | 7 | -0.792, -0.466 | | | | 8,7 | | e | | 148/150 | E,D,K,A,S,I |
| 54 | G | GLY:54:A | -0.223 | 6 | -0.466, -0.086 | | | | 7,5 | | e | | 148/150 | G,Q,N,R,S,L,D,C |
| 55 | D | ASP:55:A | -0.939 | 8 | -1.056, -0.884 | | | | 9,8 | | e | f | 148/150 | D,E,S,N,P,T,V |
| 56 | P | PRO:56:A | -0.779 | 8 | -0.928, -0.693 | | | | 8,8 | | e | f | 148/150 | P,S,L,A,F,V,W |
| 57 | C | CYS:57:A | -0.065 | 5 | -0.330, 0.121 | | | | 6,5 | | e | | 147/150 | C,N,H,L,I,Y,V,P,A |
| 58 | T | THR:58:A | -0.504 | 7 | -0.693, -0.400 | | | | 8,6 | | e | | 147/150 | T,S,R,V,I,M,A,N,P,L,Y |
| 59 | V | VAL:59:A | -0.394 | 6 | -0.585, -0.256 | | | | 7,6 | | b | | 146/150 | V,I,L,T,M,A,F |
| 60 | S | SER:60:A | -0.752 | 8 | -0.884, -0.693 | | | | 8,8 | | e | f | 147/150 | S,A,T,Y,Q,L,N,G,D,H |
| 61 | S | SER:61:A | -0.564 | 7 | -0.744, -0.466 | | | | 8,7 | | e | | 148/150 | S,P,T,D,K,R,C,L,H |
| 62 | Q | GLN:62:A | -0.475 | 7 | -0.640, -0.330 | | | | 7,6 | | e | | 148/150 | Q,P,D,E,L,S,K,F,H,R |
| 63 | L | LEU:63:A | 1.57 | 2 | 1.060, 1.497 | | | | 3,2 | | e | | 148/150 | L,D,M,A,H,R,I,Q,V,E,N,K,T,S |
| 64 | E | GLU:64:A | -0.89 | 8 | -1.014, -0.792 | | | | 9,8 | | e | f | 147/150 | E,Y,D,Q,A,X,P,S,T |
| 65 | L | LEU:65:A | -0.736 | 8 | -0.928, -0.640 | | | | 8,7 | | b | | 148/150 | L,M,F,W,S,Q,R |
| 66 | E | GLU:66:A | 0.748 | 4 | 0.389, 1.060 | | | | 4,3 | | e | | 147/150 | E,N,D,R,G,Q,K,T,Y,F,A,C |
| 67 | E | GLU:67:A | -0.985 | 9 | -1.097, -0.928 | | | | 9,8 | | e | f | 148/150 | E,D,A,T,K,Y |

| | | | | | | | | | | |
|---|---|---|---|---|---|---|---|---|---|---|
| 68 | A | ALA:68:A | -1.059 | 9 | -1.181, -1.014 | 9,9 | b | s | 149/150 | A,G,S,R,F,T |
| 69 | F | PHE:69:A | 0.308 | 4 | 0.012, 0.560 | 5,4 | b | | 149/150 | F,I,L,V,Y,S,H,M |
| 70 | R | ARG:70:A | -0.951 | 8 | -1.097, -0.884 | 9,8 | e | f | 148/150 | R,G,L,M,N,K,Y |
| 71 | L | LEU:71:A | -0.693 | 8 | -0.884, -0.585 | 8,7 | e | f | 148/150 | L,I,F,A,Q,V,C |
| 72 | Y | TYR:72:A | 0.204 | 5 | -0.086, 0.389 | 5,4 | b | | 147/150 | Y,S,F,H,T,A,C,I,L,P |
| 73 | E | GLU:73:A | 1.358 | 3 | 0.775, 1.497 | 4,2 | e | | 147/150 | E,D,S,R,C,K,H,G,N,Q,V,P,F,T,Y |
| 74 | L | LEU:74:A | 1.465 | 2 | 0.775, 1.497 | 4,2 | e | | 147/150 | L,I,R,E,Q,V,M,F,H,A,Y,K,T |
| 75 | N | ASN:75:A | -0.445 | 7 | -0.640, -0.330 | 7,6 | e | | 147/150 | N,K,T,P,E,H,R,D,S,Y,G,C |
| 76 | K | LYS:76:A | 0.829 | 4 | 0.389, 1.060 | 4,3 | e | | 148/150 | K,R,E,P,A,T,Y,H,G,N,S,Q |
| 77 | D | ASP:77:A | 0.522 | 4 | 0.121, 0.775 | 5,4 | e | | 148/150 | D,G,E,R,C,F,K,M,S,V,Q,N,T |
| 78 | S | SER:78:A | 0.122 | 5 | -0.174, 0.246 | 6,5 | e | | 148/150 | S,A,E,T,Y,D,I,P,N,C,K,R,L |
| 79 | E | GLU:79:A | -0.26 | 6 | -0.466, -0.086 | 7,5 | e | | 148/150 | E,Q,G,C,F,I,K,D,N,S,M,V |
| 80 | L | LEU:80:A | 0.088 | 5 | -0.174, 0.246 | 6,5 | b | | 148/150 | L,F,I,A,M,D,C |
| 81 | L | LEU:81:A | 0.719 | 4 | 0.389, 1.060 | 4,3 | e | | 148/150 | L,I,C,V,T,F,S,K,A,Q,N,H |
| 82 | I | ILE:82:A | 0.195 | 5 | -0.086, 0.389 | 5,4 | b | | 148/150 | I,L,V,R,M,F,N,S |
| 83 | H | HIS:83:A | -0.834 | 8 | -0.972, -0.744 | 9,8 | b | | 148/150 | H,L,G,N,Y |
| 84 | V | VAL:84:A | -0.456 | 7 | -0.640, -0.330 | 7,6 | b | | 141/150 | V,A,I,S,G,T,L,E,P,F |
| 85 | F | PHE:85:A | -0.769 | 8 | -0.928, -0.640 | 8,7 | b | | 140/150 | F,S,L,E,H,T |
| 86 | P | PRO:86:A | 0.426 | 4 | 0.012, 0.560 | 5,4 | e | | 141/150 | P,C,K,D,N,A,S,E,L,I,G |
| 87 | C | CYS:87:A | -0.185 | 6 | -0.400, 0.012 | 6,5 | e | | 140/150 | C,G,S,N,R,A,H,T,Y |
| 88 | V | VAL:88:A | 1.149 | 3 | 0.775, 1.497 | 4,2 | e | | 140/150 | V,I,L,A,T,S,E,H,P,R,Q,K,C,M |
| 89 | P | PRO:89:A | -0.954 | 8 | -1.097, -0.884 | 9,8 | e | f | 141/150 | P,E,S,L,A |
| 90 | E | GLU:90:A | 2.083 | 1 | 1.060, 2.631 | 3,1 | e | | 140/150 | E,P,S,A,V,L,Q,K,T,M,I,D,R,G |
| 91 | R | ARG:91:A | 2.027 | 2 | 1.060, 2.631 | 3,1 | e | | 136/150 | R,H,E,L,V,Q,A,K,G,I,D,S,N,C,Y |
| 92 | P | PRO:92:A | -0.738 | 8 | -0.928, -0.640 | 8,7 | e | f | 139/150 | P,A,E,S,K |
| 93 | G | GLY:93:A | -1.043 | 9 | -1.181, -0.972 | 9,9 | e | f | 139/150 | G,K,I,F |
| 94 | M | MET:94:A | 0.498 | 4 | 0.121, 0.775 | 5,4 | e | | 139/150 | M,V,K,L,D,E,T,S,I,Q,A |
| 95 | P | PRO:95:A | -0.434 | 7 | -0.640, -0.330 | 7,6 | e | | 140/150 | P,L,S,A,Q,T |
| 96 | C | CYS:96:A | -0.616 | 7 | -0.839, -0.466 | 8,7 | b | | 140/150 | C,Y,F,M,A,H,L |
| 97 | P | PRO:97:A | 1.136 | 3 | 0.560, 1.497 | 4,2 | e | | 140/150 | P,A,Q,T,H,I,E,K,V,D,C,N |
| 98 | G | GLY:98:A | -0.5 | 7 | -0.744, -0.330 | 8,6 | e | | 140/150 | G,E,T,L,Y,S,N,P |
| 99 | E | GLU:99:A | -1.059 | 9 | -1.181, -1.014 | 9,9 | b | s | 142/150 | E,F,Y,G,S |
| 100 | D | ASP:100:A | -0.576 | 7 | -0.744, -0.466 | 8,7 | e | | 142/150 | D,E,T,K,S,N,G,H,V,P |
| 101 | K | LYS:101:A | 0.405 | 4 | 0.121, 0.560 | 5,4 | e | | 146/150 | K,R,L,E,F,Q,G,S,T,P,D |
| 102 | S | SER:102:A | -0.425 | 7 | -0.585, -0.330 | 7,6 | e | | 148/150 | S,N,Y,K,H,T,R,A,L,I,F,P |
| 103 | I | ILE:103:A | -0.653 | 7 | -0.792, -0.585 | 8,7 | e | | 150/150 | I,V,M,L |
| 104 | Y | TYR:104:A | -0.787 | 8 | -0.972, -0.693 | 9,8 | e | f | 150/150 | Y,H,F,S,C,L,G |
| 105 | R | ARG:105:A | -1.012 | 9 | -1.139, -0.928 | 9,8 | e | f | 150/150 | R,H,Y,L,N,Q |
| 106 | R | ARG:106:A | -0.916 | 8 | -1.056, -0.839 | 9,8 | e | f | 150/150 | R,A,K,G,I,Y,H |
| 107 | G | GLY:107:A | -0.801 | 8 | -0.972, -0.693 | 9,8 | e | f | 150/150 | G,A,N,H,S,T,V,K |
| 108 | A | ALA:108:A | -1.129 | 9 | -1.225, -1.097 | 9,9 | e | f | 150/150 | A,S,Y,G,T |
| 109 | R | ARG:109:A | -0.908 | 8 | -1.056, -0.839 | 9,8 | e | f | 150/150 | R,G,K,H,I |
| 110 | R | ARG:110:A | -0.737 | 8 | -0.884, -0.640 | 8,7 | e | f | 150/150 | R,C,H,P,F,Q,L,K,I |
| 111 | W | TRP:111:A | -0.248 | 6 | -0.585, 0.012 | 7,5 | e | | 150/150 | W,K,P,L,F,Y,R |
| 112 | R | ARG:112:A | -0.756 | 8 | -0.928, -0.640 | 8,7 | e | f | 150/150 | R,K,T,Q,E,L,Y |
| 113 | K | LYS:113:A | -1.111 | 9 | -1.225, -1.056 | 9,9 | e | f | 150/150 | K,R,D |
| 114 | L | LEU:114:A | -0.144 | 6 | -0.400, 0.012 | 6,5 | b | | 150/150 | L,I,Q,F,M,R,Y,V,K,H |
| 115 | Y | TYR:115:A | -0.74 | 8 | -0.928, -0.640 | 8,7 | e | f | 149/150 | Y,P,H,R,L |
| 116 | C | CYS:116:A | 0.682 | 4 | 0.246, 0.775 | 5,4 | b | | 149/150 | C,K,R,Y,Q,L,H,F,N,M,I |
| 117 | A | ALA:117:A | 0.048 | 5 | -0.174, 0.246 | 6,5 | b | | 149/150 | A,I,V,L,M,W,F,N,T,P |
| 118 | N | ASN:118:A | -0.678 | 7 | -0.839, -0.585 | 8,7 | b | | 149/150 | N,L,H,S,K,I,D,M,F,Y,T,R,C |
| 119 | G | GLY:119:A | -0.907 | 8 | -1.056, -0.792 | 9,8 | e | f | 149/150 | G,T,D,Y,C |
| 120 | H | HIS:120:A | -1.152 | 9 | -1.275, -1.097 | 9,9 | b | s | 149/150 | H,C,Q,Y |
| 121 | T | THR:121:A | 0.025 | 5 | -0.256, 0.246 | 6,5 | b | | 149/150 | T,A,S,L,R,I,K,N,F,P,E,V |
| 122 | F | PHE:122:A | -0.549 | 7 | -0.744, -0.400 | 8,6 | b | | 149/150 | F,Y,H,L,V,S |
| 123 | Q | GLN:123:A | -0.63 | 7 | -0.792, -0.527 | 8,7 | e | | 149/150 | Q,V,A,L,H,T,I,R,S,E,Y |
| 124 | A | ALA:124:A | -0.822 | 8 | -0.972, -0.744 | 9,8 | e | f | 149/150 | A,G,V,S,P,K |
| 125 | K | LYS:125:A | -0.898 | 8 | -1.056, -0.792 | 9,8 | e | f | 149/150 | K,R,V,P,S |
| 126 | R | ARG:126:A | -0.71 | 8 | -0.884, -0.585 | 8,7 | e | f | 146/150 | R,C,H,Y,M,N,G,P,X,S |
| 127 | F | PHE:127:A | -0.84 | 8 | -1.014, -0.744 | 9,8 | e | f | 147/150 | F,L,P,V |
| 128 | N | ASN:128:A | -0.867 | 8 | -1.014, -0.792 | 9,8 | e | f | 146/150 | N,S,G,A,T,I |
| 129 | R | ARG:129:A | -0.548 | 7 | -0.744, -0.400 | 8,6 | e | | 147/150 | R,H,K,P,Q,T,Y,M |
| 130 | R | ARG:130:A | 0.558 | 4 | 0.246, 0.775 | 5,4 | e | | 146/150 | R,L,K,N,M,Q,T,G,H,A,C,P,V,F,X |
| 131 | A | ALA:131:A | -0.861 | 8 | -1.014, -0.792 | 9,8 | b | | 148/150 | A,T,V,I,S |
| 132 | H | HIS:132:A | 0.984 | 3 | 0.560, 1.060 | 4,3 | e | | 147/150 | H,Q,F,R,Y,N,M,D,L,S,V,T,C,E,A,F |
| 133 | C | CYS:133:A | -1.006 | 9 | -1.139, -0.928 | 9,8 | b | s | 147/150 | C,G,F,V |
| 134 | A | ALA:134:A | 0.035 | 5 | -0.256, 0.246 | 6,5 | e | | 146/150 | A,G,F,T,V,K,P,I,E,Y,H,S |
| 135 | I | ILE:135:A | 0.647 | 4 | 0.246, 0.775 | 5,4 | e | | 145/150 | I,V,Y,H,Q,F,R,L,K |
| 136 | C | CYS:136:A | -1.005 | 9 | -1.139, -0.928 | 9,8 | e | f | 144/150 | C,G,F,Y |
| 137 | T | THR:137:A | 1.022 | 3 | 0.560, 1.060 | 4,3 | e | | 144/150 | T,S,N,Q,H,R,A,G,K,E,V,L,I,M,F |
| 138 | D | ASP:138:A | -0.949 | 8 | -1.097, -0.884 | 9,8 | e | f | 144/150 | D,E,G,T,V,Y |
| 139 | R | ARG:139:A | -0.399 | 6 | -0.585, -0.256 | 7,6 | e | | 144/150 | R,C,I,G,Y,K,T,P,Q,H,D |
| 140 | I | ILE:140:A | -1.017 | 9 | -1.139, -0.972 | 9,9 | b | s | 144/150 | I,V,Y,T,L |
| 141 | W | TRP:141:A | -0.847 | 8 | -1.056, -0.744 | 9,8 | e | f | 144/150 | W,G,A |
| 142 | G | GLY:142:A | -1.041 | 9 | -1.181, -0.972 | 9,9 | e | f | 144/150 | G,R,A |
| 143 | L | LEU:143:A | -0.855 | 8 | -1.014, -0.744 | 9,8 | e | f | 144/150 | L,F,I,S |
| 144 | G | GLY:144:A | -0.932 | 8 | -1.097, -0.839 | 9,8 | e | f | 142/150 | G,A,S |
| 145 | R | ARG:145:A | -0.284 | 6 | -0.527, -0.174 | 7,6 | e | | 142/150 | R,C,Q,L,G,Y,K,S,H,I |
| 146 | Q | GLN:146:A | -0.892 | 8 | -1.014, -0.792 | 9,8 | e | f | 142/150 | Q,P,S,K,F |
| 147 | G | GLY:147:A | -0.991 | 9 | -1.139, -0.928 | 9,8 | b | s | 143/150 | G,V,A,F |
| 148 | Y | TYR:148:A | -0.008 | 5 | -0.330, 0.246 | 6,5 | b | | 143/150 | Y,F,C,V,H,L,M |
| 149 | K | LYS:149:A | -0.731 | 8 | -0.884, -0.640 | 8,7 | e | f | 143/150 | K,R,Y,N,Q,F |
| 150 | C | CYS:150:A | -1.147 | 9 | -1.275, -1.097 | 9,9 | b | s | 142/150 | C,F,X |
| 151 | I | ILE:151:A | 0.69 | 4 | 0.246, 0.775 | 5,4 | e | | 143/150 | I,T,L,M,V,N,E,C,A |
| 152 | N | ASN:152:A | -0.055 | 5 | -0.330, 0.121 | 6,5 | e | | 143/150 | N,D,H,Q,T,K,A,G,R,E,S,L |
| 153 | C | CYS:153:A | -1.215 | 9 | -1.309, -1.181 | 9,9 | e | f | 143/150 | C |
| 154 | K | LYS:154:A | -0.641 | 7 | -0.839, -0.527 | 8,7 | e | | 143/150 | K,R,P,H,Q |
| 155 | L | LEU:155:A | -0.2 | 6 | -0.466, 0.012 | 7,5 | e | | 143/150 | L,V,I,Y,M,H,P |
| 156 | L | LEU:156:A | 0.017 | 5 | -0.256, 0.246 | 6,5 | e | | 143/150 | L,S,I,C,A,M,R,T,W,V,F |
| 157 | V | VAL:157:A | -1.051 | 9 | -1.181, -1.014 | 9,9 | b | s | 143/150 | V,L,I |
| 158 | H | HIS:158:A | -1.227 | 9 | -1.309, -1.181 | 9,9 | b | s | 144/150 | H,F |
| 159 | K | LYS:159:A | -1.057 | 9 | -1.181, -1.014 | 9,9 | e | f | 143/150 | K,Q,R,F |
| 160 | K | LYS:160:A | -0.789 | 8 | -0.928, -0.693 | 8,8 | e | f | 144/150 | K,R,H,N,F,E |
| 161 | C | CYS:161:A | -1.149 | 9 | -1.275, -1.097 | 9,9 | b | s | 144/150 | C,F |
| 162 | H | HIS:162:A | -0.805 | 8 | -0.972, -0.693 | 9,8 | b | | 143/150 | H,S,R,A,Y,T,V,F |
| 163 | K | LYS:163:A | 0.164 | 5 | -0.174, 0.389 | 6,4 | e | | 143/150 | K,R,G,A,I,P,T,V,S,E,L |
| 164 | L | LEU:164:A | 0.3 | 4 | -0.086, 0.560 | 5,4 | e | | 143/150 | L,N,V,A,Y,S,K,I,C,F,T,M,Q |
| 165 | V | VAL:165:A | -0.168 | 6 | -0.400, 0.012 | 6,5 | b | | 143/150 | V,L,I,Y,F,Q,A,G,T,C |
| 166 | T | THR:166:A | 0.453 | 4 | 0.121, 0.560 | 5,4 | e | | 143/150 | T,P,R,N,Q,D,K,S,G,H,I,L |
| 167 | I | ILE:167:A | 1.606 | 2 | 1.060, 1.497 | 3,2 | e | | 143/150 | I,T,Q,R,L,K,V,H,M,D,F,N,Y,S,P |
| 168 | E | GLU:168:A | 1.697 | 2 | 1.060, 2.631 | 3,1 | e | | 143/150 | E,S,T,V,P,A,K,D,G,Y,I,R,L,Q,H |
| 169 | C | CYS:169:A | -1.002 | 9 | -1.139, -0.928 | 9,8 | e | f | 142/150 | C,S,W,Y |
| 170 | G | GLY:170:A | 0.839 | 4 | 0.389, 1.060 | 4,3 | e | | 120/150 | G,Q,R,T,P,S,K,H,A,N,E,D |
| 171 | R | ARG:171:A | 0.146 | 5 | -0.174, 0.389 | 6,4 | e | | 88/150 | R,Q,E,H,N,L,D,S,T,K,A |
| 172 | H | HIS:172:A | -0.085 | 5 | -0.400, 0.121 | 6,5 | e | | 53/150 | H,R,Q,N,P,Y |
| 173 | S | SER:173:A | -0.704 | 8 | -1.014, -0.527 | 9,7 | e | f | 16/150 | S,T |

*Below the confidence cut-off - The calculations for this site were performed on less than 6 non-gaped homologue sequences,
or the confidence interval for the estimated score is equal to- or larger than- 4 color grades.

### Table S3

phosphorylation sites identified for aPKCι in the MaxQuant database (MaxQB)

| Sequence | Type | Residue | Position | Identifications | Best Score | Best Score Diff | Best Localization Probability | Regions | Conserved In |
|---|---|---|---|---|---|---|---|---|---|
| PGDTTST(ph)FCGTPNY | Phospho (STY) | T | 11 | 2 | 121.23 | 6.63664 | 0.808082 | Required for interaction with RAB2, Regulatory domain | |
| TVAGGGS(ph)GDHSHQV | Phospho (STY) | S | 19 | 1 | 205.12 | 25.6809 | 0.997304 | Required for interaction with RAB2, Regulatory domain | |
| RRWRKLY(ph)CANGHTF | Phospho (STY) | Y | 136 | 1 | 57.175 | 43.9897 | 0.99996 | Regulatory domain | |
| NEPVQLT(ph)PDDDDIV | Phospho (STY) | T | 564 | 1 | 364.21 | 108.536 | 1 | AGC-kinase C-terminal | |

### Table S4

Primers and constructs used in this study

| | | |
|---|---|---|
| cloning primers | | backbone |
| PKCi_FlagReg_FW | GTGGCGGCCGCTCGAGATGGACTACAAAGACGATGACGACAAGCCGACCCAGAGGGAC | pcDNA3.1 |
| PKCi_Reg_RV | AAACGGGCCCTCTAGATTAATCCTGAAGACCTAGACTGGATG | pcDNA3.2 |
| Myc_aPKCi_KD_FW | GTGGCGGCCGCTCGAGATGGAGCAGAAGCTGATCTCAGAG | pcDNA3.1 |
| aPKC_RV | AAACGGGCCCTCTAGATCAGACACATTCTTCTGCAGACATC | pcDNA3.1 |
| aPKC_Reg_TEV_FW | TGGAGCCATCCGCAATTTG | pcDNA3.1 |
| aPKC_Reg_TEV_RV | TGACTGGAAGTACAGGTTTTCATGC | pcDNA3.1 |
| aPKC_Kin_Strep_FW | CTGTACTTCCAGTCATCTTTGCCACAGGAACCAGTG | pcDNA3.1 |
| aPKC_Kin_Strep_RV | TTGCGGATGGCTCCAGACACATTCTTCTGCAGACATC | pcDNA3.1 |
| mutagenesis primers | | |
| PKCi_A129E_FW | TAGAGGTGAACGCCGCTGGAGAAAGCTT | |
| PKCi_A129E_RV | CGGCGTTCACCTCTACGGTAGATGGATTTATCT | |
| PKCi_Y136F_FW | AAAGCTTTTTTGTGCCAATGGCCACACTTTCC | |
| PKCi_Y136F_RV | GCACAAAAAAGCTTTCTCCAGCGGCG | |
| PKCi_Y136E_FW | AAAGCTTGAATGTGCCAATGGCCACACT | |
| PKCi_Y136E_RV | GCACATTCAAGCTTTCTCCAGCGGCGT | |
| PKCi_C1_R147A_FW | AGCCAAGGCTTTCAACAGGCGTGCTCACTGTGC | |
| PKCi_C1_R147A_RV | TTGAAAGCCTTGGCTTGGAAAGTGTGGCC | |
| PKCi_C1_R150/151A_FW | TCAACGCGGCTGCTCACTGTGCCATCTGC | |
| PKCi_C1_R150/151A_RV | GAGCAGCCGCGTTGAAACGCTTGGCTTGG | |
| PKCi_C1_RIWGL_FW | CGCAATAGCGGGAGATGGACGCCAAGGATATAAGTGC | |
| PKCi_C1_RIWGL_RV | TCTCCCGCTATTGCGTCTGTGCAGATGGCACAGTG | |
| PKCi_PS4xRA_FW | CTGCAGGTGCAGCCGCCTGGAGAAAGCTTTATTGTGCC | |
| PKCi_PS4xRA_RV | CGGCTGCACCTGCAGCGTAGATGGATTTATCTTCTCCTGG | |
